# PEG-mCherry interactions beyond classical macromolecular crowding

**DOI:** 10.1101/2024.05.07.592799

**Authors:** Liam Haas-Neill, Khalil Joron, Eitan Lerner, Sarah Rauscher

## Abstract

The dense cellular environment influences bio-macromolecular structure, dynamics, interactions and function. Despite advancements in understanding protein-crowder interactions, predicting their precise effects on protein structure and function remains challenging. Here, we elucidate the effects of PEG-induced crowding on the fluorescent protein mCherry using molecular dynamics simulations and fluorescence-based experiments. We identify and characterize specific PEG-induced structural and dynamical changes in mCherry. Importantly, we find interactions in which PEG molecules wrap around specific surface-exposed residues in a binding mode previously observed in protein crystal structures. Fluorescence correlation spectroscopy experiments capture PEG-induced changes, including aggregation, suggesting a potential role for the specific PEG-mCherry interactions identified in simulations. Additionally, mCherry fluorescence lifetimes are influenced by PEG and not by the bulkier crowder dextran or by another linear polymer, polyvinyl alcohol, highlighting the importance of crowder-protein soft interactions. This work augments our understanding of macromolecular crowding effects on protein structure and dynamics.

## Introduction

Macromolecular crowding can influence the structural equilibrium of proteins ^1–3^, but the precise nature of the change is difficult to predict. Since cellular environments are highly crowded, understanding the effects of macromolecular crowding on protein structure is crucial to understanding protein function in a cellular context. The result of crowding on protein structure and stability can be understood as the combination of two contributing effects: excluded volume effects (i.e., “hard interactions”) and weak, isotropic and non-specific interactions (i.e., “soft interactions,” “chemical interactions,” or “quinary interactions”) ^4–11^. Crowders take up space around a protein, inhibiting it from accessing states that would otherwise occupy the obstructed space, giving rise to the excluded volume effect ^12–14^. Soft interactions refer to any interaction not captured by treating crowders as hard spheres ^4^ and are determined by the chemical nature of the crowding agent ^4;8;15^. Soft interactions can enhance the excluded volume effect through additional repulsion or counterbalance it with attractive interactions ^4;6;8;15^. Importantly, crowder-macromolecule interactions are thought to be mainly non-specific ^16^. The isotropic interactions of crowders with macromolecules contribute to microviscosity ^17;18^, which is also modulated by depletion effects ^19;20^.

The influence of crowders on the conformational landscape of a protein is more prominent for intrinsically disordered proteins (IDPs) than for structured proteins ^21–23^. Since IDPs have many accessible states with varying degrees of compactness, the excluded volume effect strongly influences their conformational ensembles ^21^. Within the context of the conformational ensemble of an IDP, extended conformations are less favoured energetically, and hence, more compact conformations become relatively more stable. Computational models that account for the excluded volume effect but not for soft interactions have shown that IDP ensembles become more compact with increasing crowder concentration ^5;24^. Soft interactions also affect IDPs strongly. Since IDPs have shallow, rugged energy landscapes ^25^, the perturbation of many soft interactions has a more pronounced effect when compared to the effects of soft interaction perturbations on the deeply funnelled energy landscapes of folded proteins ^22;26^. Since soft interactions can be repulsive or attractive ^4;6;8^, their influence on protein structure is varied, with some IDPs exhibiting compaction and others exhibiting expansion as a function of increasing crowder size or concentration ^7;27^.

The varied crowding-induced structural effects highlight the complexity of the relationship between protein structure and crowding and the challenge in predicting these effects ^1^. Simulations with explicit crowders offer a powerful tool to disentangle the contributions of hard and soft interactions to the observed effects of crowding on the structural equilibrium and dynamics of proteins. Computational studies of proteins in crowded environments have only recently become feasible due to advances in computational power and efficiency ^1;2;8;9;22;23;28–36.^ Given the ability to model the environment of choice, molecular dynamics (MD) simulations are useful in probing the structural properties of a protein in various environments ^1;34;37;38^. Recent all-atom MD studies have investigated changes to the structure of disordered regions due to crowding and have used atomistic insights to understand the structural changes induced by crowders ^1;8;9;22;36^.

The effect of crowders on folded proteins has been studied extensively, with a major focus on the relative stability of the folded state. ^2;30;39–46^ Changes to the folded state structure itself are often not studied, challenging to probe, or considered negligible. This is likely because crowders are commonly treated as non-interacting rigid bodies acting solely via excluded volume effects, destabilizing less-folded conformations but not influencing the structure of the well-folded conformation per se. Many well-folded proteins function through transitions between conformational states; hence, crowding can also affect their function via changes to their conformational equilibrium ^1;3^. For instance, crowding influences the function of the bacterial transcription initiation complex, which exerts its function through conformational changes ^47;48^. However, studies reporting crowding-induced changes in folded proteins with a predominant conformation indicate, at most, a modest change in structure ^10;22;36;43;44;46;49^. In that respect, fluorescent proteins (FPs) provide a recent example of crowding having a significant and observable influence on the function (i.e., its fluorescence) of a highly stable folded-state protein via subtle effects on its structure and dynamics ^50^.

FPs are naturally occurring protein-caged fluorescent molecules that autocatalytically form a chromophore from three consecutive amino acids ^51^. Due to the self-sufficient nature of their fluorescence and their high fluorescence quantum yield (i.e., relative to the chromophore out of the context of the protein), they are widely used for visualizing gene expression and protein localization in cells ^51;52^. The FPs used today result from generations of directed evolution and protein design to improve a range of fluorescence properties ^53^. Monomeric red FPs (mRFPs), such as mCherry, are among the most popular because their red emission can be distinguished from autofluorescence in most cells and from the fluorescence of another popular FP, green FP (GFP). Additionally, owing to their monomeric nature, they can be genetically fused with other proteins without undergoing dimerization or tetramerization, which could affect their function ^53;54^. The structure of known FPs consists of a *β*-barrel fold encompassing a chromophore-containing *α*-helix ^55^ (Figure 1a). The *α*-helix is threaded through the center of the barrel ^55^, and additional helices and coils cross over the top and bottom of the cylindrical barrel ^56–58^.

**Figure 1:**
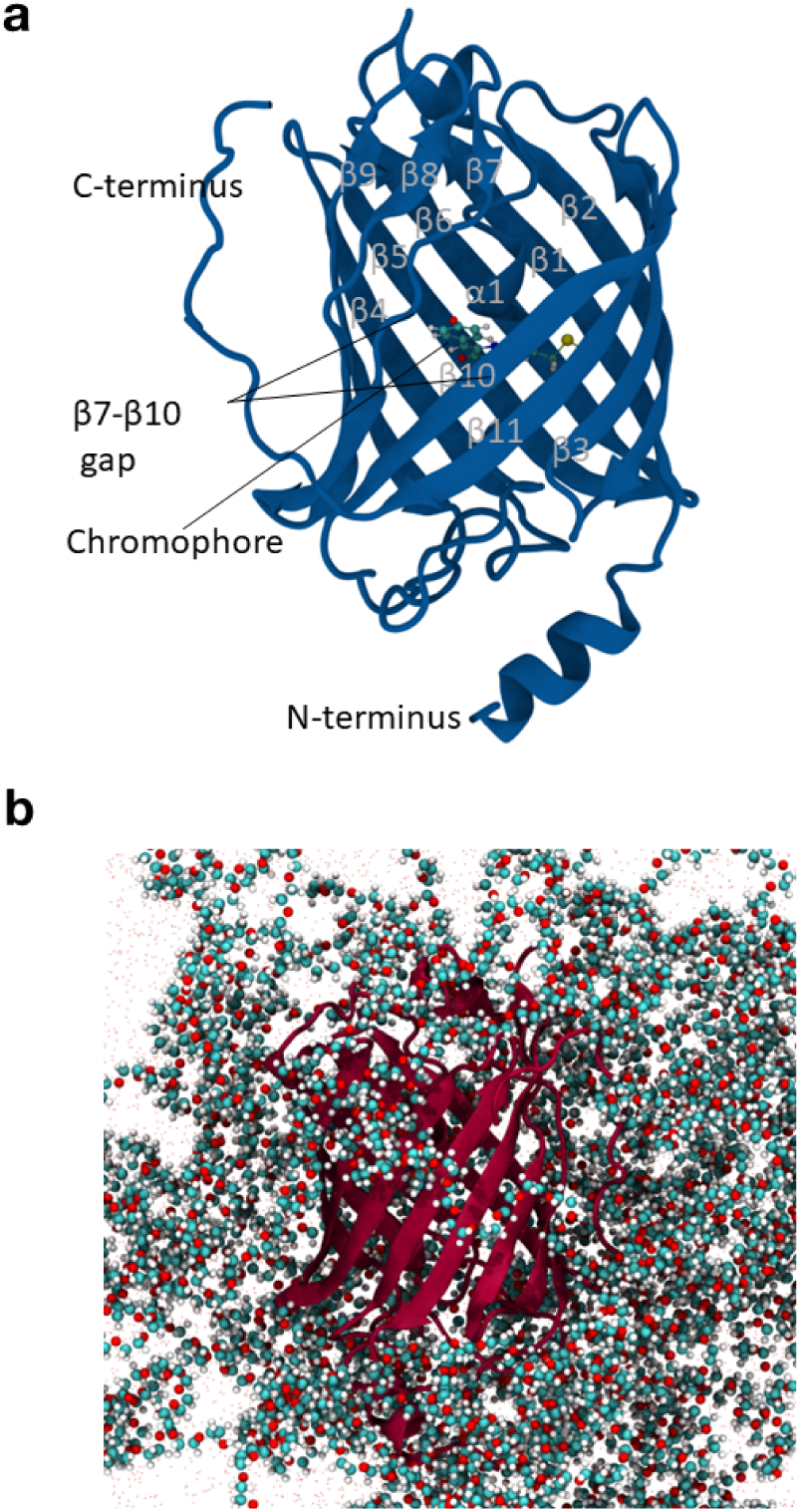
mCherry Structure. (a) The structure of mCherry with structural elements labeled. The same protein construct was used in simulations and FCS experiments. The structure shown is from a simulation of this construct. (b) Representative frame from a simulation showing mCherry in the presence of PEG. The structure of mCherry is shown in cartoon representation, and the chromophore and PEG are shown in ball and stick representation.

There is a close link between the structure of FPs and their fluorescence properties ^55;59–61^. For instance, the sequences of some FP spectral variants can differ by only a single amino acid in the *β*-barrel ^55^. The fluorescence lifetime, *τ*, of an FP is influenced by the chemical characteristics of the chromophore’s immediate environment, as well as the rigidity the protein environment imposes on the chromophore. The chemical and physical characteristics of the chromophore’s environment are, in turn, affected by the structure and dynamics of the protein, as well as by the solvent environment ^59–61^. Recent experimental work has shown that the fluorescence lifetime of various FPs can decrease in the presence of elevated concentrations of crowders ^50^. *τ* was observed to monotonically decrease with increasing crowder concentrations above 30% fractional volume occupancy (FVO) with no corresponding spectral changes. This trend was observed for small molecular crowders or osmolytes, such as trehalose, and different sizes of polyethylene glycol (PEG). The lifetime reduction effect was purely due to crowding and was most pronounced for mCherry, among all the FPs tested. Furthermore, this study showed that slight expansion of the chromophore’s environment, or the chromophore pocket, in PEG-crowded conditions reduced constraints on the chromophore. Time-resolved fluorescence anisotropy measurements supported this finding, as they showed that the chromophore of mCherry gains rotational degrees of freedom relative to the barrel fold in the presence of ≥ 30 % FVO of PEG. Based on these findings, the expansion of the chromophore pocket was suggested to be the reason for the reduced fluorescence lifetime of mCherry in the presence of PEG ^50^.

Here, we performed all-atom MD simulations of mCherry in water and in the presence of PEG (Figure 1) to gain molecular-level insights into PEG-protein interactions and how these interactions affect the protein’s structure. Our simulations reveal that PEG induces a conformational change in mCherry that subtly influences the local structure of the *β*-barrel, specifically involving residues important for fluorescence. We study the nature of the interactions between PEG and mCherry and find that PEG forms specific interactions with the protein, rather than behaving as an ideal, inert crowder. The results of fluorescence correlation spectroscopy (FCS) measurements of mCherry in the presence of varying concentrations of PEG align with the findings from simulations. Additionally, fluorescence lifetime measurements show that flexible polymeric crowders, such as PEG, permit maximizing crowder-protein soft interactions, non-specific and specific alike, compared to bulkier crowders, such as dextran. We also compare the effects of PEG on fluorescence lifetime changes in mCherry to those in the presence of another linear polymeric agent, polyvinyl alcohol (PVA), which has different chemical features than PEG. Our results are consistent with specific PEG-mCherry interactions and their unique effects on mCherry’s fluorescence characteristics. Altogether, the results highlight how both the rigidity and the chemical properties of a crowder play an important role in its soft interactions with macromolecules, which can have a significant impact on the structure and function of a macromolecule in general and of a fluorescent protein in particular.

## Materials and Methods

### System Setup & Simulation Protocol

The crystal structure of mCherry was taken from the Protein Data Bank (PDB ID 2H5Q) ^55^. Missing atoms were added with the CHARMM-GUI PDB-reader tool ^62^. Missing residues at the termini were added to match the UNIPROT sequence (UNIPROT ID: X5DSL3), and a six-residue histidine tag was added to the C-terminus with UCSF Chimera ^63^. This sequence was used to match the construct used for the *in vitro* experiments in Joron *et al.* (2023) ^50^. The initial structures of all residues added to the crystal structure were modeled with the MODELLER loop refinement tool ^64^.

Refer to Figure S1a for the sequence of the mCherry construct. Five structures were generated using MODELLER, and the one with the lowest DOPE-HR score ^65^ (high-resolution version of the Discrete Optimized Protein Energy) was used as the initial structure for simulations.

All simulations were carried out as part of the study of Joron *et al.*, ^50^ and the simulation protocol is briefly summarized here. Simulations were performed using GROMACS version 2022.2^66^.

The mCherry anionic chromophore parameters ^67^ were converted to GROMACS format using the charmm2gmx python script with modifications for Python 3 compatibility. The CHARMM36m force field ^68^ was used with the CHARMM-modified TIP3P water model ^69^. The Nand C-termini were simulated in the charged state, and all titratable residues were simulated in their standard states for pH 7. For the simulations of the protein in water, the protein was placed in a triclinic box with 1 nm distance to all box edges to obtain the system’s initial configuration. Periodic boundary conditions were used. Water molecules ^69^ were added to the simulation box, and neutralizing Na^+^ and Cl^-^ ions were added at a concentration of 139.7 mM. The system had a total of 38,774 atoms. Energy minimization was performed using the steepest descent method. The v-rescale thermostat ^70^ was used for all simulations, and velocities were generated from a Maxwell distribution. The system was equilibrated for 10 ns in the NVT ensemble with 1,000 kJ/mol position restraints on all protein heavy atoms, followed by a 10 ns constant pressure simulation using the Berendsen barostat ^71^ without changing the position restraints. The Parrinello-Rahman barostat ^72^ was used for all subsequent simulations. Next, the system was equilibrated for 20 ns in the NPT ensemble without changing the position restraints. Lastly, the system was simulated without any position restraints for 100 ns. The final frame of this simulation was used as the starting configuration for five independent production simulations, which were run for 2 *µ*s in the NPT ensemble.

To generate conformations of PEG in water, we simulated a system consisting of 200 PEG molecules in water. This system, consisting of 200 molecules of 28 consecutive ethylene glycol units (total of 199 atoms and molecular weight 1.25 kDa, i.e., equivalent to PEG 1250) and 6,184 water molecules, was set up using the CHARMM-GUI ^62^ polymer-builder tool. The energy of the system was minimized using the steepest descent method. The system was equilibrated in the NVT ensemble with the Nośe-Hoover thermostat ^73^ for 250 ps, followed by a 1 ns equilibration simulation in the NPT ensemble using the Nośe-Hoover thermostat and the Parrinello-Rahman barostat ^72^. Next, a 100 ns equilibration simulation in the NPT ensemble using the v-rescale ^70^ thermostat and the Parrinello-Rahman barostat ^72^ was carried out. A new simulation box containing the pre-equilibrated structure of mCherry from the simulations in water was used for the initial setup of the PEG system. All water molecules and ions were removed. A random PEG molecule was extracted from the final frame of the PEG in water simulation and inserted into this new simulation box. The GROMACS tool gmx insert-molecules ^66^ was used to insert the PEG molecules. Once gmx insert-molecules was unable to insert any more copies of the PEG molecule into the system, a new random PEG molecule with a different conformation was extracted from the PEG in water simulation, and gmx insert-molecules was used again to insert it. This process was repeated until choosing a new PEG conformation did not allow for any new PEG molecules to be added to the simulation box, for a total of six iterations and 62 PEG molecules. Water molecules were then added to the system along with neutralizing Na^+^ and Cl^-^ ions at a concentration of 139.7 mM for a final PEG concentration of ∼42% FVO. The PEG system contained a total of 36,353 atoms. The energy of the system was minimized using the steepest descent method. Subsequent steps in the simulation protocol were the same as described above for the no-PEG system.

### Analysis of Simulation Trajectories

The first quarter of each simulation was delineated as an equilibration period based on the radius of gyration (R*_g_*), number of hydrogen bonds, and backbone root-mean-square deviation (RMSD) time series reaching a plateau (Figure S3). After the equilibration period, the RMSD between the structure of mCherry in each trajectory ranged from 3.2 to 8.3 Å, indicating that the structures were considerably dissimilar. The equilibration period was then omitted from all subsequent analyses. All coordinate-based trajectory analysis was performed using the MDAnalysis module version 2.2.0^74^ for Python 3, including R*_g_*, atomic distances, root-mean-square fluctuation (RMSF), dihedral angles, and contacts. The secondary structure was analyzed with the DSSP ^75^ function in the MDTraj ^76^ module for Python 3. Solvent accessible surface area (SASA) was computed using the Shrake-Rupley algorithm ^77^ with a probe radius of 1.4 Å and 960 sphere points in the MDTraj module for Python 3. Hydrogen bonds were computed using MDAnalysis with a 3.5 Å distance cutoff between donor and acceptor and a donor-hydrogen-acceptor angle cutoff of 150°. Two residues were considered to be in contact based on a 3.5 Å cutoff between any pair of atoms. Spatial distributions were computed with the gmx spatial tool in GROMACS ^66^ and visualized with VMD ^78^. Uncertainty is reported as the standard error of the mean, with each independent trajectory being treated as an independent measurement. The Jensen-Shannon distance ^79^ and the corresponding backbone torsion angles were computed using the PENSA ^80^ package for Python 3 with 10 bins. In the analysis of the *β*7-*β*10 gap, the autocorrelation was computed in Python 3, and fit using a sum of two exponentials with the form given in Equation 1 since the data was not well fit by a single exponential (Figure S4). Average timescales were computed as the weighted average to correctly account for the weighting of each exponential (see Equation 2).

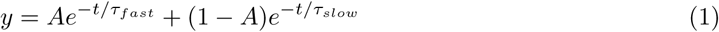

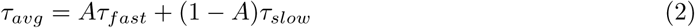

The chromophore pocket residues are the set of protein residues that form contacts with the chromophore during the simulations. Residues forming contacts in less than 0.33% of frames were excluded. This cutoff was chosen such that highly transient interactions were excluded. The pocket volume was defined as the convex hull volume of all C*α* atoms of the chromophore pocket residues. The convex hull was determined with the Open3D ^81^ module for Python 3 after extracting the atomic coordinates using MDAnalysis ^74^. gmx mindist from GROMACS version 2020.1^82^ was used to check the minimum distance between periodic images in the trajectories. Several instances of contact between period images occurred (about 3% of frames, Figure S5a-b). The correlation of these contacts with mean chromophore pocket volume in each trajectory was checked and no correlation was found (Figure S5c). The probability distributions of multiple structural properties were computed using (1) using all simulation frames, and (2) excluding frames in which mCherry had contacts with periodic images (Figure S6). All pairs of distributions were the same, regardless of whether these frames were included or excluded, and so all simulation frames were used for analysis.

### Fluorescence Correlation Spectroscopy Measurements

We performed fluorescence lifetime measurements and FCS measurements using a confocal-based microscopy setup (ISS^TM^, Champaign, IL, USA) assembled on top of an Olympus IX73 inverted microscope stand (Olympus, Tokyo, Japan). We used 488 nm pulsed picosecond diode laser (QuixX^®^ 488-60 PS, Omicron, USA; pulse width of 200 ps FWHM, operating at 80 MHz repetition rate and 160 *µ*W, measured at the back aperture of the objective lens) for exciting mCherry. The laser beam passes through a polarization-maintaining optical fiber (P1-405BPM-FC-Custom, with specifications similar to those of PM-S405-XP, Thorlabs, Newton, NJ, USA). After passing through a collimating lens (AC080-016-A-ML, Thorlabs), the beam is further shaped by a linear polarizer (DPM-100-VIS, Meadowlark Optics, Frederick, CO, USA) and a halfwave plate (WPMP2-20(OD)-BB 550 nm, Karl Lambrecht Corp., Chicago, IL, USA). A major dichroic mirror with high reflectivity at 532 and 640 nm (ZT532/640rpc, Chroma, Bellows Falls, Vermont, USA), reflects the light to the optical path through galvo-scanning mirrors (6215H XY, Novanta Corp., Boston, MA, USA) and a scan lens (30 mm Dia. x 50mm FL, VIS-NIR Coated, Achromatic Lens, Edmund Optics, Barrington, NJ, USA; both used to acquire the scanned image), and then into the side port of the microscope body through its tube lens, positioning it at the back aperture of a high numerical aperture (NA) super apochromatic objective (UPLSAPO60XW, 60X, NA=1.2, water immersion, Olympus), which focuses the light onto a small effective excitation volume, positioned within the sample chamber (*µ*-Slide 18 Well high Glass Bottom, Ibidi, Gräfelfing, GmbH). Scattered light returns in the excitation path and a fraction of it is imaged on a CCD camera, using Airy ring pattern visualization for z-positioning. Fluorescence from the sample is collected through the same objective, is transmitted through the major dichroic mirror and is focused with an achromatic lens (25mm Dia. x 100mm FL, VIS-NIR Coated, Edmund Optics) onto a 100 *µ*m diameter pinhole (variable pinhole, motorized, tunable from 20 *µ*m to 1 mm, custom made by ISS^TM^), and then re-collimated with another achromatic lens (AC254-060-A, Thorlabs). Fluorescence is then further cleaned using a 615/24 nm single-band bandpass filter (FF01-615/24-25, Semrock, Rochester, NY, USA). Fluorescence was collected using one hybrid PMT, routed to a TCSPC card (SPC 150N, Becker & Hickl, GmbH) with a TAC window of 12.5 ns. Data acquisition is performed using the VistaVision software (version 4.2.424, 64-bit, ISS^TM^) in the time-tagged time-resolved (TTTR) file format.

FCS data acquisitions were performed for 5-30 minutes on ∼1 nM mCherry in PBS buffer containing varying concentrations of PEG 6,000. Next, the fluorescence autocorrelation function was calculated from the acquired fluorescence trajectories, as previously shown ^83^. The fluorescence autocorrelation functions were fitted to a model that includes up to two translational 3D diffusion processes and one exponential rapid dynamics process (see Equation 3).

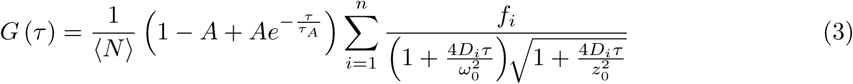

where *τ* is a lag time, *G*(*τ*) is the autocorrelation of the data at different lag times, *< N =* is the mean number of fluorescent molecules in the effective excitation volume, *D_i_* is the translational diffusion coefficient of a component *i* and *f_i_* is the fraction of that component out of *n* diffusional components of two at most. *ω*_0_ and *z*_0_ are the width and height of the effective excitation volume, respectively. *τ_A_*and *A* are the relaxation time and the equilibrium fraction, respectively, of a dim state of a fluorescence fluctuation process between bright and dim states (see Equation 4).

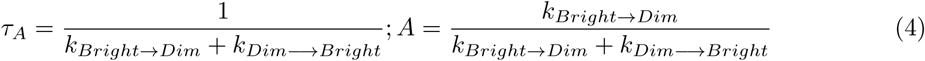

We report the fluorescence traces, their autocorrelation functions, and their model-fitting results. We also report the values of the mean hydrodynamic radii calculated out of the best-fit results of the translational diffusion process as well as the best-fit values of the rapid dynamics process. The mean hydrodynamic radii were calculated by using the Stokes-Einstein equation, assuming spherical particles (see Equation 5), and by using the dimensions of the effective excitation volume, which were estimated from best-fit results to the fluorescence autocorrelation function of a small chromophore with a diffusion coefficient of ∼320 *µm*^2^*/s*, ATTO 532^84^.

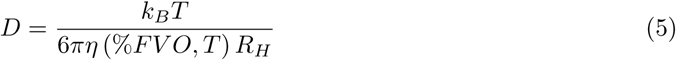

where *k_B_* is the Boltzmann constant, *T* is the absolute temperature, *η* (%*FV O, T*) is the bulk viscosity of the solution of buffer at a given crowder concentration (i.e., % FVO) at a given temperature and *R_H_* is the hydrodynamic radius.

Additionally, fluorescence decays were acquired for 100 nM mCherry (addgene plasmid mCherry: #29722) in PBS buffer in the absence and presence of varying concentrations of PEG 6,000 or 10,000 (Sigma-aldrich #81253 and #92897, respectively), as well as dextran 5,000 or 10,000 (Sigma-aldrich #31404 and #D9260, respectively), or PVA 10,000 (9,000 - 11,000) (Sigma-aldrich #360627). The mean fluorescence lifetimes were calculated from the best-fit results of fitting the fluorescence decays to a convolution of a sum of exponentials function with the recorded impulse response function (IRF; see Equation 6).

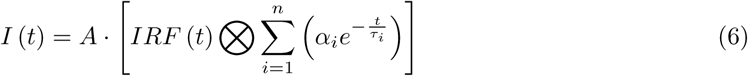

where *t* is the time after a pulsed excitation, better known as the time-correlated single photon counting (TCSPC) time, *I* (*t*) is the model of the fluorescence decay, *IRF* (*t*) is the IRF, *n* is the number of exponential decay components, which is two at most, *τ_i_* and *α_i_* are the fluorescence lifetime and the relative amplitude of a given exponential component, respectively, and *A* is a proportionality factor. The mean fluorescence lifetime, *τ*, was then calculated as follows (see Equation 7).

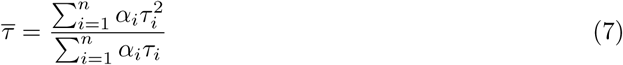

The propagated errors, *δf* (*p_i_*), for the values of calculated parameters, *f* (*p_i_*) (e.g., *R_H_*, *τ*), were calculated as follows (see Equation 8).

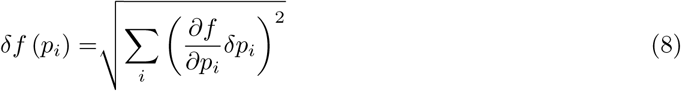

where *δp_i_*are the fitting errors for the best-fit parameter values *p_i_*. Analyses of the FCS data and the representation of all results were performed using the OriginLab Origin software version 2022b.

## Results

We performed simulations of mCherry in water and in the presence of 42% FVO PEG (Figure 1b; see Table S1 for a list of simulation systems). The description of the results is organized as follows. First, we describe how PEG affects the structured *β*-barrel region of the protein. Then, we describe the influence of PEG on the structural dynamics of the flexible *β*7-*β*10 gap, followed by a discussion of the influence of PEG crowding on the disordered N- and C-terminal regions of the mCherry construct. We also discuss specific interactions between mCherry and PEG, including a binding mode in which PEG molecules wrap around solvent-exposed side chains of specific residues. Then, we discuss the results of FCS experiments, which suggest a potential relation between the slowdown of conformational dynamics observed in the MD simulations and a slowdown in fluorescence fluctuations. Additionally, FCS results reveal mCherry aggregation in the presence of both PEG and dextran, suggesting one outcome of specific PEG-mCherry interactions. Finally, while we have previously shown that PEG leads to a reduction in the fluorescence lifetime of mCherry ^50^, here, we show that dextran, a bulkier crowder, does not affect the fluorescence lifetime. Further comparisons based on FCS measurements of mCherry in the presence of PEG and dextran suggest that while dextran acts as expected for a classical excluded volume crowder, PEG interacts and influences mCherry aggregation and structural dynamics.

### PEG Influences Local Structure in the Ordered Region of mCherry

Our previous study ^50^ demonstrated that the fluorescence lifetime of mCherry decreases as a function of increasing PEG concentration above 30% FVO. We found that this decrease in fluorescence lifetime can be explained by a slight expansion of the *β*-barrel near the chromophore, which increases the degrees of freedom available to the chromophore inside the barrel, ultimately leading to more excited-state fluorescence quenching and, hence, to shorter fluorescence lifetime ^50^. Here, we aim to provide a complete description of how the structure and dynamics of mCherry are altered in an environment crowded by PEG. First, we analyzed the structure of the pocket that surrounds the chromophore (Figure 2a; see Table S2 for a list of residues constituting the chromophore pocket). We find that the radius of gyration (*R_g_*) of the pocket is larger on average in the presence of PEG crowders (Figure 2b). This result is consistent with our previous findings using the pocket volume as a metric of the space accessible to the chromophore ^50^. Next, we take a broader view of how PEG crowding affects the structured region of the protein.

**Figure 2:**
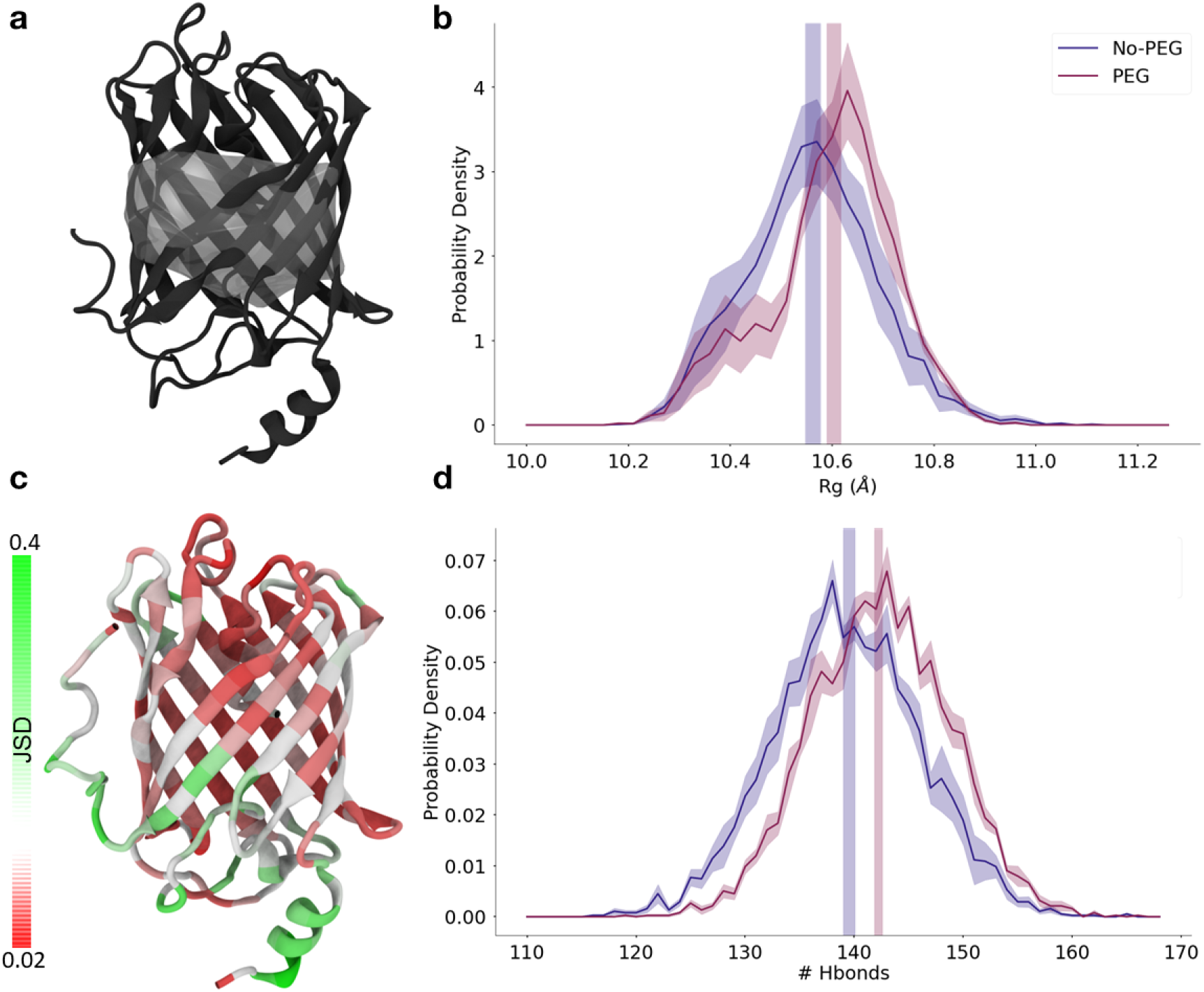
PEG causes subtle changes in the local structure of the ordered region of mCherry. (a) The 3D structure of mCherry is shown in the new cartoon representation. The chromophore pocket is shown in grey in a translucent space-filling representation. (b) The *Rg* distributions of the C*α* atoms of the chromophore pocket residues are shown with and without PEG (plum and navy, respectively). (c) The Jensen-Shannon distance (JSD) of backbone dihedral angle distributions between the PEG and no-PEG systems is shown mapped onto the structure. Red indicates smaller JSD, green indicates larger JSD, and white indicates intermediate values. (d) The distributions of the number of intra-protein hydrogen bonds in the *β*-barrel of mCherry (excluding tails, residues 13-220) are shown with and without PEG (plum and navy, respectively). In (b) and (d), the shaded areas indicate the standard error of the mean obtained by treating each trajectory as independent.

To fully characterize the influence of PEG on the structure of mCherry, we also investigate structural changes distant from the chromophore. While we observe significant changes in the size of the chromophore pocket (Figure 2b), there is no difference in the global dimensions of the ordered part of the protein due to PEG, as measured by R*_g_* (Figure S7b). We also computed the number of hydrogen bonds in the *β*-barrel (Figure 2d), finding that the number of intra-barrel hydrogen bonds is significantly higher in the presence of PEG. This observation can be explained by the fact that PEG molecules are less available to form hydrogen bonds than water. When PEG obstructs water from the protein’s surface, fewer hydrogen bonds are formed between the protein and water, leaving some residues free to form additional protein-protein hydrogen bonds.

We wanted to investigate whether the structural changes due to PEG occur isotropically across the protein surface or if some regions are more perturbed than others. For this purpose, we utilized the Jensen-Shannon distance (JSD) ^79;80^ to compare the backbone dihedral distributions between the systems with and without PEG. The fact that the JSD varies greatly between residues (Figure 2c) indicates that the influence of PEG on the structure of the ordered region of mCherry is non-uniform. The JSD is highest for the disordered N- and C-terminal regions, which indicates that the influence of PEG is attenuated for the structured region overall compared to the disordered regions. These findings are consistent with an analysis of contacts between pairs of residues (Figure S8), which also reveals a non-uniform change in structure across the *β*-barrel and an increased perturbation of contacts in the disordered tails compared to minimal changes to the structure of the *β*-barrel.

A particularly important region of the *β*-barrel for the fluorescence properties of mCherry is the *β*7-*β*10 gap (Figure 3a). Like other monomeric derivatives of DsRed, mCherry has a gap in the *β*- barrel between the *β*7 and *β*10 strands. This gap is a vestige of the missing tetrameric interactions disrupted by mutations introduced to design a monomeric FP ^85^. Previous studies have shown that the *β*7-*β*10 gap undergoes structural transitions between closed and open states ^60^, allowing molecular oxygen to diffuse into the barrel and affect the photostability of the chromophore ^54;85^.

**Figure 3:**
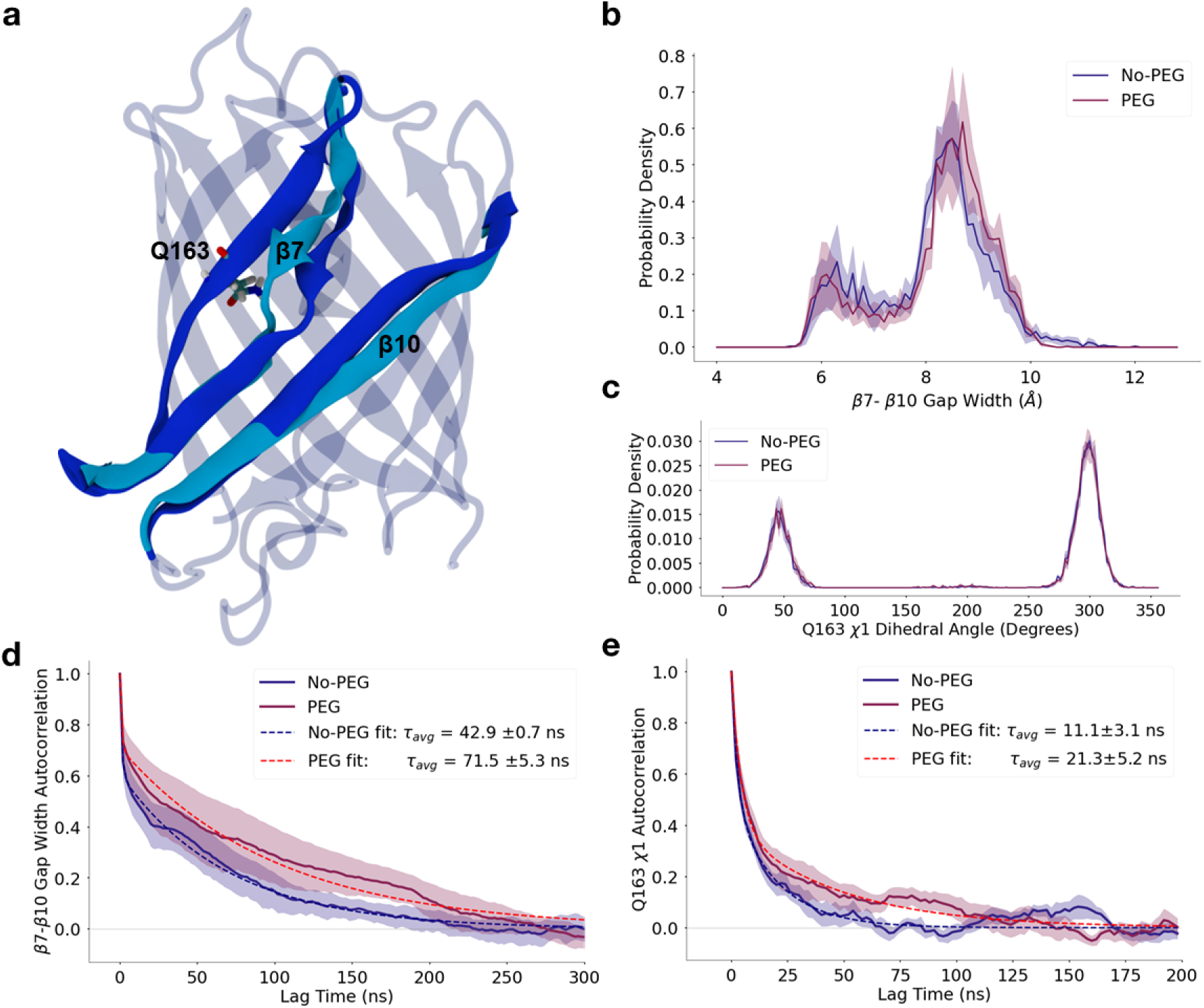
PEG affects the transition dynamics of the. *β***7-***β***10 gap, but not its structure or its structural equilibrium.** (a) Representative 3D structures of the open (light blue) and closed (dark blue) states of the *β*7-*β*10 gap. Q163, located on *β*8, is shown in licorice representation. (b) The distribution of *β*7-*β*10 gap width for both systems. The gap width is computed as the distance between the center of mass of the C*α* atoms of three residues on *β*7 (E144, A145, S146) and three residues on *β*10 (I197, K198, L199). (c) The distribution of the *χ*1 dihedral angle of residue Q163 for both systems. (d-e) The autocorrelation of the *β*7-*β*10 gap width and Q163 dihedral angle, respectively, for both the PEG and no-PEG systems. The sum of two exponentials fit (Equation 1) to each autocorrelation is shown with the weighted average of the characteristic timescales (Equation 2), *τav*, provided in the legend (see Methods for details). The fits for the PEG and no-PEG systems are shown in red and blue dotted lines, respectively. For the plots in (b), (c), (d), and (e), the results for the PEG system and no-PEG systems are shown in plum and blue, respectively. Shaded regions indicate the standard error of the mean obtained by treating each trajectory as independent.

To determine the width of the *β*7-*β*10 gap, we measured the distance between three residues on *β*7 (E144, A145, S146) and three residues on *β*10 (I197, K198, L199). Consistent with previous studies ^54;60^, we observe that the *β*7-*β*10 gap undergoes transitions between two highly populated states, as seen in Figure 3b. The state where the gap distance is small corresponds to the closed state, and the state where the gap distance is large corresponds to the open state. We observe that the gap is more likely to be in the open state than the closed state, and the presence of PEG does not affect the populations of these states. Notably, the *β*7-*β*10 gap provides a clear example of how similar the protein structure is with and without PEG. Mukherjee *et al.* also observed two states of the gap, albeit with slightly different populations than those observed here ^60^. This difference in state populations may be due to differences in force field choice, simulation system, or conformational sampling (i.e., 2 *µ*s vs. 20 *µ*s in the present work, see Table S1).

Our results indicate that the presence of PEG does not affect the populations of the conformational states of the *β*7-*β*10 gap. However, there are other possible pathways for molecular oxygen to diffuse to the chromophore. Previous studies have shown that the side chain of residue 163 can block the diffusion of molecular oxygen to the chromophore through either the *β*8-*β*7 gap, which opens up when *β*7 is close to *β*10^54^, or through cavities inside the protein, depending on the orientation of its side chain ^85^. To determine if PEG alters the side chain orientation of Q163, we analyzed the *χ*_1_, *χ*_2_, and *χ*_3_ side chain dihedral distributions of this residue. We observe no difference in the rotameric state populations due to PEG (Figure 3c, Figure S9), which suggests that there would be no difference in oxygen access to the chromophore through the *β*-barrel in the presence of PEG. Therefore, these results imply that the decrease in fluorescence lifetime observed in experiment ^50^ cannot be attributed to differences in the structure of Q163, the *β*7-*β*10 gap, or the diffusion of molecular oxygen.

In order to investigate the relationship between the state of the *β*7-*β*10 gap and the volume of the chromophore pocket, we decomposed the chromophore pocket volume distribution into separate distributions for each state. To do this, we fitted a sum of two Gaussian functions to the *β*7-*β*10 gap width distributions for each system and used the local minimum between the two peaks as a cutoff point to define the open and closed states (Figure S10a). We then assigned simulation frames to either the open or closed state based on this cutoff. To ensure the robustness of this method, we repeated this state assignment using an alternate method for defining the width of the *β*7-*β*10 gap (the same method used in Mukherjee *et al.* ^60^, Figure S10b). Both methods result in a bimodal distribution with the open state being more populated than the closed state.

We then computed the probability distributions of the chromophore pocket volume separately for the open and closed states of the *β*7-*β*10 gap (Figure S10c,d). These distributions show that when the *β*7-*β*10 gap is in the closed state, the pocket volume is unaffected by PEG, whereas in the open state, the pocket volume is larger in PEG than in water. This difference likely arises because of the rigidity conferred by the *β*-strand interactions in the closed state; the hydrogen bonds between *β*-strands prevent the structural perturbation induced by PEG. Joint distributions show moderate correlation between the *β*7-*β*10 gap width and the pocket volume (Figure S11a-d). Although the populations of the open and closed states of the *β*7-*β*10 gap are not affected by PEG, these states are directly related to the changes in the chromophore pocket volume, which, in turn, affects the fluorescence lifetime.

Overall, we observe subtle local changes to the structure of the ordered region of the protein, including the chromophore pocket. Previous studies on the influence of PEG crowding on structured proteins have described similarly subtle effects ^10;22^. One simulation study found that an ordered protein slightly expands upon crowding ^22^, similar to the expansion of the chromophore pocket observed in our simulations. An experimental study has shown that PEG can induce local disorder in structured regions of proteins and that these subtle structural changes can influence protein function ^10^. Our findings align with these works - we observe that PEG crowding exerts small yet non-negligible influences on certain parts of the *β*-barrel of mCherry. The slight expansion of the *β*-barrel in the chromophore pocket provides a plausible mechanism for the reduced fluorescence lifetime observed for mCherry in the presence of crowders ^50^.

### PEG Slows Down The Conformational Dynamics of mCherry

Because PEG-induced crowding causes slight changes to the size of the chromophore pocket and the structure of the *β*-barrel (Figure 2b, d), we wanted to investigate if the structural dynamics of mCherry are also altered. To this end, we analyzed transitions in the width of the *β*7-*β*10 gap. We computed the autocorrelation of the *β*7-*β*10 gap width, which was best modelled using a sum of two exponentials (Equation 1), to determine the characteristic relaxation times (Figure 3d and Table S3; see Methods for details). In the presence of PEG, the relaxation timescale is slower by a factor of ∼1.7, relative to the absence of PEG. We then performed the same autocorrelation analysis on the side chain fluctuations of Q163 (Figure 3e, Table S3). The average relaxation timescale for the side chain fluctuations is nearly two times slower in PEG than without PEG. While the Q163 *χ*_1_ rotameric state populations are identical in the crowded and uncrowded systems, the time required for transitions from one state to the other differs. Similarly, the timescale of *β*7-*β*10 gap fluctuations is significantly slower in PEG. Together, these results indicate that PEG crowding slows the rates of important conformational changes involving the *β*7-*β*10 gap, even though it does not change the populations of the conformational states.

Previous works that studied the influence of crowders on macromolecule structural dynamics have shown that crowders can influence translational and rotational diffusion timescales of proteins ^1;8;9;16;36;86–90^ and small molecules ^91^. Most studies have found that crowders lead to longer diffusion timescales ^1;8;9;16;36;86–90^, although some have suggested that faster diffusion is possible in specific cases ^36;91^. In general, the increase in diffusion timescales is thought to result from the restriction of free space due to the presence of crowder molecules rather than due to soft interactions with the crowder ^8^. Given that conformational fluctuations often require diffusion of parts of the protein through the solvent, our observations of slowed conformational transition rates are consistent with these previous findings of slowed diffusion due to the presence of crowders and their induced viscosity. Moreover, multiple pieces of evidence indicate that crowding can influence conformational transition rates ^36;47;48;92^. For instance, Ostrowska *et al.* found that crowders can slow protein side chain fluctuations by 20-50% for crowder concentrations of ∼15% FVO ^36^. Our results show a slowing of conformational fluctuations involving the *β*7-*β*10 gap by between 66% and 100% (Table S3). These findings are consistent with those of Ostrowska *et al.*, particularly given that the extent to which dynamics are slowed is expected to scale with crowder concentration. Because we used a significantly higher crowder concentration (42% FVO compared to 15% FVO used by Ostrowska *et al.*), the fact that we observe a more significant slowdown of conformational fluctuations is to be expected.

### PEG Enhances Helicity and Overall Compaction of the IDRs

Since the influence of PEG on the structure of intrinsically disordered regions (IDRs) has been extensively studied ^9;23;36^, it is useful to study the disordered regions in mCherry to test if PEG similarly perturbs its IDRs. The crystal structure of mCherry (PDB:2H5Q ^55^) indicates that the N- and C- terminal tails each contain an eight-residue disordered segment that is not resolved ^55^. These eight N-terminal residues are also not resolved in another crystal structure of mCherry (PDB:6YLM^93^). However, the C-terminal residues are resolved in that structure, possibly because crystal contacts stabilize these residues in a particular conformation. Our mCherry construct, which was selected to match the construct used in Joron *et al.*, contains these disordered segments, as well as a six-residue histidine tag at the C-terminus. To assess the flexibility of these regions in simulation, we computed the root-mean-square fluctuation (RMSF) of the protein (Figure 4b). We observe significant flexibility in both of these IDRs in the absence of crowders. In addition, the disordered regions at both the N- and C-termini are longer than the crystal structures indicate; the disordered regions in our construct each constitute 17 residues (Figures 4a-b, Figure S1b, and Figure S2). The increased flexibility we observe for mCherry in solution may be due to the absence of crystal contacts that stabilize the tails in the crystal structures. An additional contribution to the difference in flexibility in solution vs. crystal may be the conditions of the crystallization buffer, which included 30% PEG ^55^. Including the His-tag, which is not present in the crystallized construct, may also play a role. To investigate the influence that PEG has on the two IDRs, we compare the structural ensembles of these regions with and without PEG crowding. While the RMSF of the structured region is similar between the crowded and uncrowded systems, the RMSF of the N-terminal tail is significantly lower in the presence of PEG (Figure 4b, Figure S12a). The RMSF of the C-terminal tail is also slightly lower in the presence of PEG (Figure 4b, Figure S12b). The excluded volume effect is expected to lead to reduced flexibility of IDPs upon crowding since reduced access to extended states implies less interconversion between compact and extended states. Thus, the reduced flexibility of the N- and C-terminal tails in the presence of PEG is consistent with expectations based on the effect of excluded volume.

**Figure 4:**
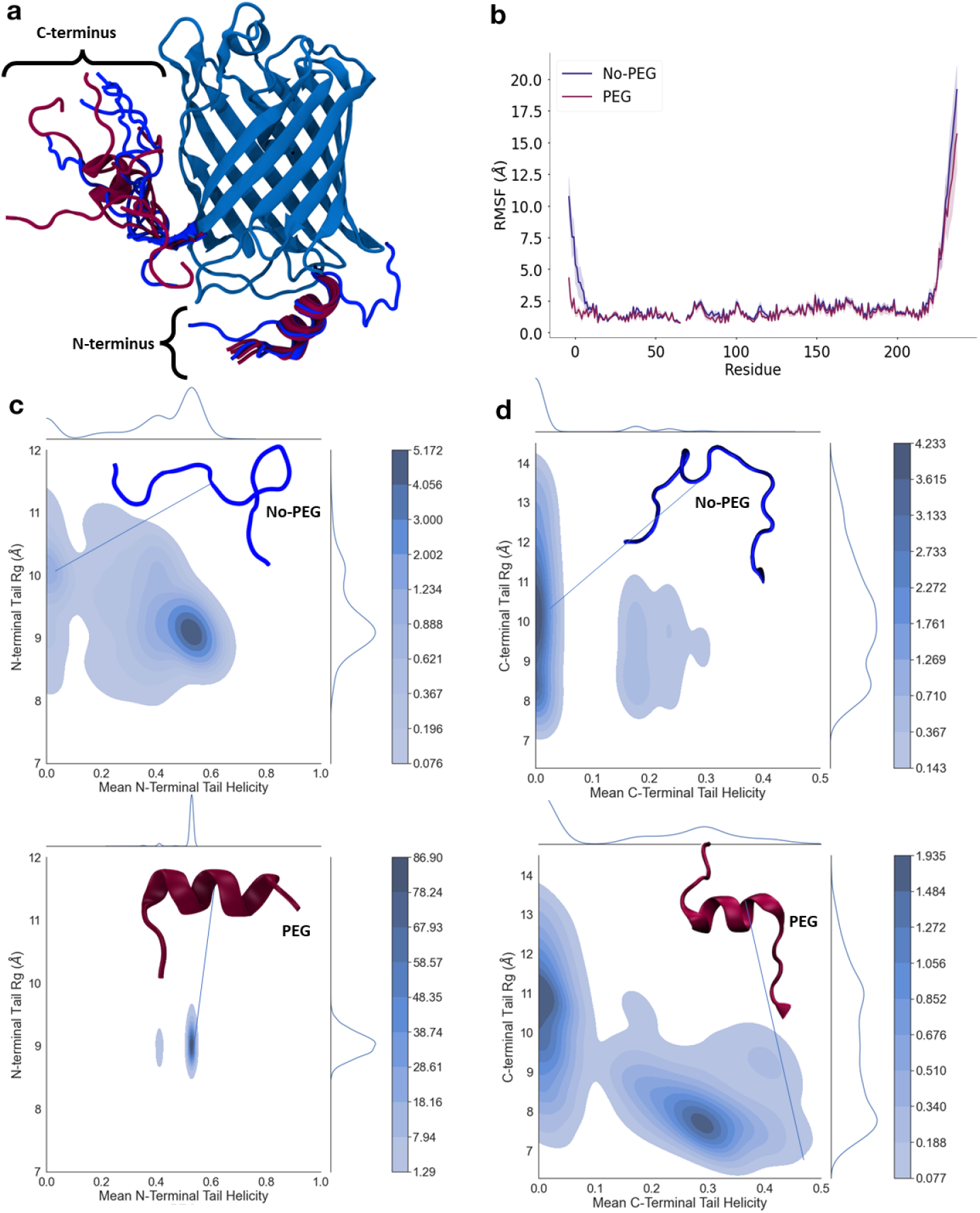
The influence of PEG crowding on the disordered regions of the mCherry construct. (a) Representative structures of the N- and C-termini for the PEG (plum) and no-PEG (blue) systems. Five structures are shown for each system. (b) Root-mean-square fluctuation (RMSF) for the PEG system (plum) and the no-PEG system (blue). Shaded regions indicate the standard error of the mean, treating each system as an independent observation. There is a gap in the RMSF profile because the chromophore comprises three residues but is only assigned one RMSF value. (c-d) 2D histograms of the mean helicity vs R*g* of the N- and C- terminal regions. The top histograms are for the no-PEG system, and the bottom histograms are for the PEG system. Representative structures from selected regions in the phase space are shown. 2D histograms for the N-terminal tail and C-terminal tail are shown in (c) and (d), respectively. 1D histograms for the R*g* and helicity are provided on each plot. The kernel density estimate option in the seaborn jointplot function was used for histogram smoothing.

To better understand the structural origin of the reduced flexibility in PEG, we computed the helicity and *R_g_*of the N- and C- terminal tails. This analysis, represented as a 2D histogram of helicity and *R_g_* (Figure 4c-d, Figure S7a and c), reveals that both IDRs have two states: one with high helicity and low R*_g_*, and one with higher R*_g_* and no helicity. When PEG is present, there is a significant increase in the relative population of the states with high helical propensity and low R*_g_*. Much of the N-terminal tail remains helical in the presence of PEG and exhibits significantly reduced disorder. Most crystal structures of mCherry indicate that the N-terminal tail is not helical and is missing the first 8-10 residues ^55;93–98^. However, one crystal structure (PDB ID: 5FHV ^99^) indicates helicity in the N-terminal tail, including residues 3-8^99^. Since we observe elevated levels of helicity in the N-terminal tail in the presence of PEG compared to water (Figure S2a), we speculate that the helicity observed in the N-terminal tail in this crystal structure could be in part due to the presence of 40% FVO PEG in the crystallization buffer.

Previous studies have shown that crowding can lead to either expansion or compaction for IDPs ^7;27^. However, IDPs most often experience compaction ^9^ since the excluded volume effect destabilizes extended conformational states. Several studies have reported that crowding can stabilize helical structures in IDPs ^7;9;22^, which has also been attributed to the excluded volume effect ^9^. Using single-molecule FRET experiments, Stringer *et al.* established that an IDR experiences compaction with large PEG crowders and expansion with small PEG crowders ^27^. The relative size of PEG crowders is determined by a threshold at *P* = ^√^*N* ^27^, where P is the degree of polymerization of the crowder, and N is the degree of polymerization of the IDR. For *P >* ^√^*N*, polymers are considered large, and for *P <* ^√^*N*, polymers are considered small. In the current study, the PEG crowders used are ethylene glycol polymers consisting of 28 repeating subunits. The disordered N- and C-terminal tails of the mCherry construct each consist of 17 residues. Thus, the PEG crowders in our simulations are large relative to the size of the IDRs. The IDR compaction we observe is, therefore, consistent with the work of Stringer *et al.* ^27^ and provides additional support for this model of PEG’s influence on proteins with IDRs and IDPs.

Furthermore, previous studies have found that IDPs are more affected by crowders than folded proteins ^21;22^. To assess the relative effect of PEG-induced crowding on the disordered and structured regions of mCherry, we computed a difference map of the intra-protein contact propensities in the presence and absence of PEG (Figure S8). The largest differences in intra-protein contacts occur within the N- and C- terminal tails, which are characterized by an increase in contacts when PEG is present. Based on the differences in contacts, the structural ensemble of the IDRs is more strongly perturbed by the crowded environment than the *β*-barrel. This result is corroborated by the fact that the JSD in the IDRs is much larger than in the *β*-barrel (Figure 2c). Our results are qualitatively consistent with earlier observations that disordered proteins experience more pronounced structural changes upon crowding than structured proteins ^21;22^, and build on these previous works to show that there is a larger relative effect of crowding on IDRs compared to a structured region within the same protein.

Overall, the influence of PEG-induced crowding on the disordered regions of our mCherry construct is consistent with prior studies of IDPs and IDRs in crowded environments. Our simulations reveal that the disordered regions of mCherry display increased helical propensity, become more compact, and experience reduced flexibility in the presence of PEG. These changes, although the most pronounced we observe for this system, occur far from the chromophore. Importantly, they are not correlated with changes in the volume of the chromophore pocket (Figure S11e-h), which strongly suggests that the impacts of PEG on the structured and disordered regions of mCherry are occurring independently. Moreover, these results suggest that the pronounced effects of PEG on the IDR ensembles are not related to the observed changes in fluorescence lifetime ^50^.

### PEG Wraps Around Specific Positively Charged Residues

Though PEG is often used as a mimetic for protein crowders because it is putatively inert, previous studies have reported attractive or repulsive PEG-protein interactions ^27;100^. In this section, we investigate in detail the interactions between PEG and mCherry in our simulations. First, we computed the distribution of the number of PEG molecules that mCherry is in contact with throughout the simulations (Figure S13). This distribution shows that mCherry is in contact with 27 PEG molecules on average and never fewer than 14. To determine if PEG preferentially binds specific residues, we computed the probability that each residue is in contact with PEG during the simulation (Figure S14a,b). As expected, residues that protrude into the solvent are more likely to interact with PEG. In particular, residues in the N- and C-terminal tails and the loops are significantly more likely to be in contact with PEG than residues in the *β*-barrel (Figure S14a,b). Residues whose side chains point into the *β*-barrel have significantly less contact with PEG than their neighbours whose side chains point outwards into the solvent (Figure S14a). We also find that residues with higher solvent-accessible surface area (SASA) are more likely to interact with PEG (Figure S15), which is expected since enhanced solvent accessibility implies higher PEG accessibility. These results are consistent with PEG acting as an inert crowder.

However, we wanted to determine if there are any preferential PEG-protein interactions. Therefore, we analyzed the PEG contact probabilities by residue type (Figure 5b), finding that positively charged residues form contacts with PEG more frequently than other residue types. To investigate the nature of these preferential interactions more deeply, we computed the number of PEG atoms in contact with residues of each type (Figure S16a). We find that negatively charged residues are in contact with fewer PEG atoms on average compared to positively charged residues. Overall, these results are consistent with Ostrowska *et al.*, who found that the oxygen atoms of PEG interact less frequently with negatively charged residues and that most residues within the protein interact with PEG to some extent, with buried residues being less likely to form contact with PEG ^36^. Another simulation study found differences in PEG binding affinity between different residue types ^101^. The consistency between our results and those in previous works suggests that a preference for interaction with positive residues and against interaction with negatively charged residues may be a general feature of PEG-protein interactions regardless of structure.

**Figure 5:**
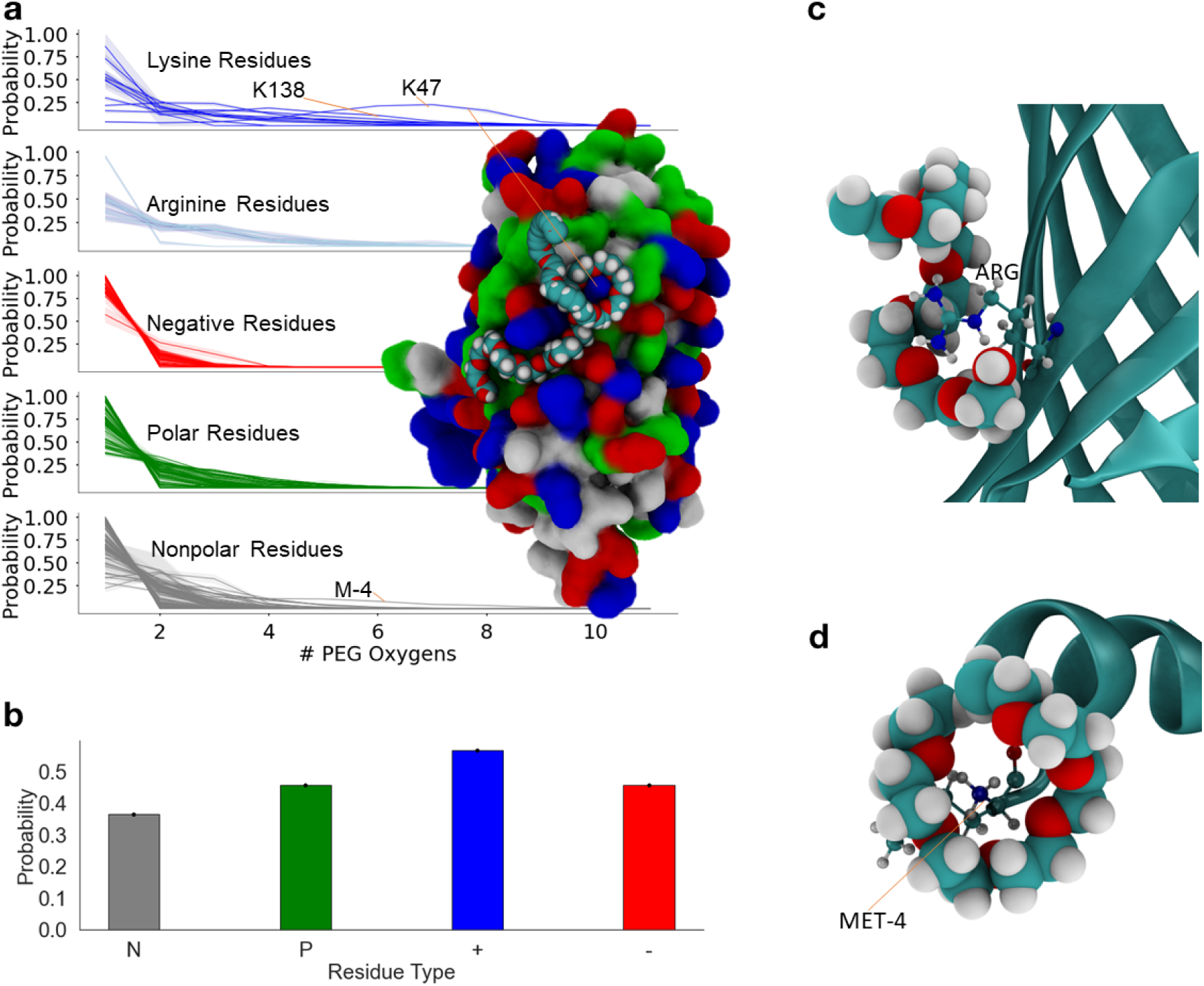
PEG-protein interactions depend on residue type. (a) The probability that each residue is in contact with a certain number of PEG oxygen atoms, broken down by residue type; frames where PEG is not in contact with each residue are excluded from this calculation. When a PEG molecule wraps around a residue, more of its oxygen atoms are in contact with that residue. Two lysine residues that have a particularly high likelihood of forming contacts with *≥* 6 PEG oxygen atoms are indicated. A representative structure with a PEG molecule wrapping around residue K47 on mCherry is shown. PEG is shown in van der Waals representation with carbon, oxygen, and hydrogen atoms shown in cyan, red, and white, respectively. mCherry is shown in surface representation with residues coloured by type. (b) The probability that residues of each type are in contact with PEG at any given simulation frame. This is computed as the probability for each residue to be in contact with PEG, which is then averaged over all residues of each type. Uncertainty is indicated with black bars. In panels (a) and (b), positively charged residues are shown in blue, negatively charged residues are shown in red, polar residues are shown in green, and non-polar residues are shown in grey. (c) A representative structure of PEG partially wrapped around an arginine residue. (d) A representative structure of PEG wrapping around the NH3 group of the N-terminal residue, MET-4. In panels (c) and (d), protein residues are shown in a ball and stick representation, and PEG is shown in a van der Waals representation.

To further investigate why positively charged residues preferentially interact with PEG, we looked at the binding modes of PEG to mCherry. In all trajectories, we observe a binding mode in which PEG molecules wrap around the solvent-exposed side chains of specific residues (most notably, residues M-4, K47, and K138; Figure 5a). To quantify the prevalence of this binding mode, we computed the number of PEG oxygen atoms in contact with each protein residue (Figure 5a, Figure S16b). This analysis demonstrates that the wrapping binding mode is particularly prevalent for K47 and, to a lesser extent for K138 (Figure 5a, Figure S16b). The prevalence of this binding mode can be seen in the 3D spatial distribution of PEG oxygen atoms around the protein (Figure 6a). The most prevalent PEG-binding “hotspot” occurs at K47 and several residues surrounding it due to the PEG wrapping at this site (Figure 6a-c, Figure S16b). Time-series analysis of the number of PEG oxygen atoms in contact with K47 (Figure S17) shows that K47 remains in contact with a PEG molecule for timescales as long as 1 *µ*s. When PEG wraps around K47, its oxygen atoms are oriented toward the positively charged side chain and its CH_2_-groups are oriented toward the surrounding residues (representative structure shown in Figure 5a). The local environment of K47 (Figure 6b) makes it an optimal environment for the PEG wrapping mode. A movie showing a representative instance of PEG wrapping around the side chain of residue K47 is provided as supporting information (Video 1).

**Figure 6:**
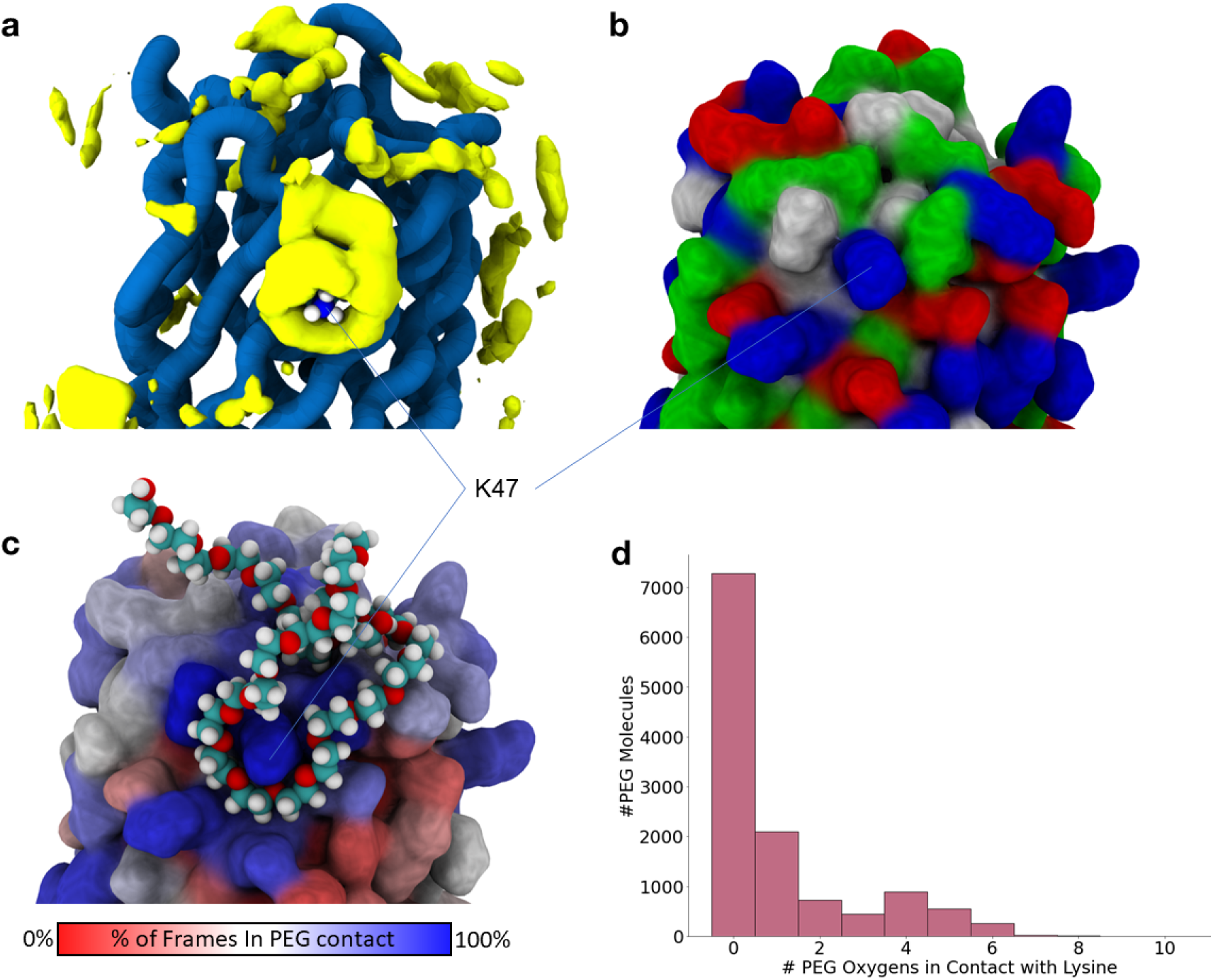
PEG wrapping at K47 is a PEG contact hotspot. (a) The 3D spatial density distribution of PEG, averaged over all simulations with PEG, is shown in yellow. Residue K47 is shown in a van der Waals representation, and mCherry is shown in a tube representation. (b) mCherry is shown in surface representation with residues coloured by residue type. Positively charged residues are shown in blue, negatively charged residues are shown in red, polar residues are shown in green, and non-polar residues are shown in grey. (c) mCherry is shown in surface representation. Residues are coloured by the percentage of simulation frames in which they are in contact with PEG. Dark red indicates 0%, and dark blue indicates 100%. A PEG molecule is shown in van der Waals representation. (d) A histogram showing the number of PEG oxygen atoms that are in contact with lysine residues in the PDB. This analysis accounts for all PEG molecules in crystal structures that contain PEG molecules of 3-10 subunits.

To determine if the PEG wrapping could be linked to the changes in the structured region leading to the experimentally observed decrease in fluorescence lifetime, we plotted a joint distribution of the chromophore pocket volume and PEG wrapping around K47 (Figure S11k). We also computed the joint distribution between the PEG wrapping of K47 and the width of the *β*7-*β*10 gap (Figure S11i,j). We find no correlation between these observables. This result strongly suggests that PEG wrapping of residue K47 is not directly responsible for the observed structural changes in the chromophore pocket.

The binding mode in which PEG wraps around lysine residues observed in our simulations has been previously observed in the crystal structure of an anti-PEG antibody ^102^, in a crystal structure of mCherry ^99^ (Figure S18), as well as in several other studies ^103–106^. The crystal structure of an anti-PEG antibody also shows lysine residues binding to a crown ether ^102^, and cyclic ethers are known to bind to cations ^107^. Multiple crystal structures include PEG molecules coordinating positively charged lysine, arginine, histidine, and potassium ions ^103^. Cation-PEG binding is used as a technique for the synthesis of pseudo-crown ethers from PEG ^108^. Another MD simulation study observed PEG wrapping around potassium, but not sodium, ions ^109^.

Our data indicate that while PEG wrapping does occur to some extent for arginine residues, it is more prominent for lysine residues (Figure 5a, Figure S16b). Lysine has a higher probability of being in contact with 6-8 oxygen atoms in PEG, whereas it is more probable for arginine to be in contact with 3-4 oxygen atoms in PEG. This difference likely stems from the geometry of the side chains of lysine and arginine. The positive charge on arginine is delocalized between its two NH_2_ groups. In simulations, the nitrogen atoms have a negative partial charge. For this reason, they act to interrupt the PEG-side chain interactions that lead to the wrapping mode (Figure 5c), resulting in contact with only 3-4 oxygen atoms in PEG at a given time. Additionally, the positive charge on arginine is more spread out in space compared to lysine. Lysine has a single NH_3_ group in its side chain. In simulations, its three hydrogen atoms carry a partial positive charge. For this reason, it is preferable for PEG to wrap around the side chain with its oxygen atoms pointing inward toward the hydrogen atoms. A compact and rotationally symmetric positive charge distribution is also present on the N-terminal NH_3_ group of MET-4, which explains why we observe the PEG-wrapping mode on that residue as well (Figure 5a,d).

It is important to note that the prevalence of PEG wrapping varies significantly even between different lysine residues on the protein’s surface (Figure 5a, Figure S19). This variation indicates that the wrapping mode depends on both the type of side chain, as well as the local environment, and therefore on protein sequence and structure. Since PEG wraps around lysine with its oxygen atoms pointing in and its CH_2_ groups pointing out, we hypothesized that PEG wrapping would be more prevalent for lysine residues surrounded by non-polar residues. We present several examples of lysine residues with varying amounts of PEG wrapping and their local environments in Figure S20. From our data, we observe that lysine residues that are in proximity to other positively charged side chains have reduced PEG wrapping (Figure S20d). Lysine residues that protrude less than other nearby surface residues also experience less PEG wrapping (Figure S20c). An enhancement of PEG wrapping is seen for lysine residues that protrude into the solvent or are near non-polar side chains (Figure 6a-c, Figure S20a,b). We show all examples of PEG wrapping of the PEG antibody crystal structure ^102^ in Figure S21 and note that all of the lysine residues that experience PEG wrapping in that protein are distant from other positively charged residues, consistent with our findings for lysine residues in mCherry. These observations align with a study that identified a similar trend, where PEG wrapping occurs for lysine residues surrounded by hydrophobic patches on the surface of the protein *λ*_6_*_−_*_85_ ^104^.

To gain insight into the prevalence of PEG wrapping in an experimental context, we analyzed a sample of protein crystal structures. First, we identified all protein structures in the PDB that contain PEG molecules consisting of 3-10 ethylene glycol subunits. We then computed the number of PEG oxygen atoms in contact with lysine residues in each of those structures. Of the 4,403 structures containing sufficiently long PEG molecules, 12% (539 structures) have lysine residues in contact with at least 5 oxygen atoms in PEG, and 20% (877 structures) have lysine residues in contact with at least 4 oxygen atoms in PEG (the complete list of these structures is provided as Supporting Material, see Data Availability). The number of oxygen atoms in contact with a lysine residue for each PEG molecule in the dataset is shown in Figure 6d. A little over half of the PEG molecules in the dataset are not in contact with any lysine residue. However, a peak with increased frequency occurs for 4-5 oxygen atoms, coinciding with the PEG wrapping binding mode. We show 3D structures for nine instances of PEG wrapping in Figure S22. Since PEG wrapping can occur with a varying number of PEG oxygen atoms, there is no precise definition of whether or not PEG wrapping is occurring for a given lysine residue. If we use 4 oxygen atoms in PEG as a minimum number to define the wrapping mode, our analysis of protein crystal structures suggests that PEG wrapping is present in at least 20% of PEG-containing crystal structures. This corresponds to around 0.4% of all structures in the PDB. Taken together, this analysis of protein crystal structures demonstrates that PEG wrapping around lysine residues is a common interaction mode between PEG and protein that is not unique to mCherry.

While the interaction between crowders and proteins is typically attributed to volume exclusion and non-specific interactions ^5–11;24^, the PEG wrapping mode is more reminiscent of a specific interaction since it occurs with high probability, and only for certain residues (Figure 5a, Figure S16b). Since PEG does not act isotropically, it cannot be considered a classical excluded volume crowder nor a simple viscogen. Furthermore, relatively strong attractive interactions between PEG and protein have been observed using force experiments, which indicated that PEG is not an inert, simple polymeric crowder, but rather it can bind proteins ^15^. Other studies ^27;110^ have also found specific interactions between PEG and proteins, suggesting that these interactions are likely a common feature of PEG-protein interactions.

### PEG Inner and Terminal Subunits Interact with the Protein Differently

Next, we investigated if there is a difference in interactions between the hydroxyl terminating groups and the oxyethylene interior groups of the PEG molecule with the protein. Differential interaction between PEG interior and end groups has previously been suggested ^111^. For all PEG molecules in contact with the protein, we computed the probability of contact formation for each PEG subunit (Figure S23). The interior of each PEG molecule tends to interact less frequently with the protein, while the PEG termini tend to interact more frequently. This difference in internal versus terminal PEG-protein contact could explain the differences in IDP expansion and collapse as a function of PEG length observed experimentally by Stringer *et al.* ^27^. Although we do not have data on varying PEG lengths from our simulations, we speculate that while longer PEG molecules occupy more space, they may be less likely to interact with proteins per monomeric unit since they have more interior groups. As a result, the effect of PEG-induced crowding on proteins with IDRs and IDPs would have a larger contribution from the excluded volume effect and a smaller relative contribution from soft interactions as PEG length increases. This dependence of interactions on PEG length might explain why longer PEG crowders cause chain compaction, while smaller PEG crowders lead to chain expansion ^27^. Nevertheless, we note that all experimentally tested PEG molecules (PEG 3,500, 6,000, and 10,000) are significantly longer than the PEG molecules used in simulation (PEG 1,250).

### Experimental Corroboration Using Fluorescence Correlation Spectroscopy

The simulation results report on interactions of PEG with mCherry (Figure 5), an increase in the pocket volume, as well as effects on the structural dynamics of the *β*7-*β*10 gap and the side chain of Q163, with relaxation times in the tens of nanoseconds (ns) (Figure 3). Other structural fluctuations of mCherry are likely also to be slowed in the presence of PEG. If such ns structural fluctuations affect the chromophore, leading to fluorescence fluctuations, and if these fluorescence fluctuations occur on timescales as rapid as hundreds of ns, then such dynamics should appear as a dynamic process with exponential relaxation in fluorescence autocorrelation functions of FCS measurements.

Additionally, we hypothesize that PEG crowding above a certain PEG concentration, as well as specific, stable PEG interactions with mCherry, could mediate bringing multiple mCherry proteins together to form aggregates. This aggregation, in turn, should influence the translational diffusion beyond the obvious effects of PEG viscosity on mCherry diffusion. To test these predictions, we performed FCS measurements of mCherry in the presence of increasing PEG concentration (i.e., 0, 20, 30, 40, and 50 % FVO PEG 6,000). We then fit the resulting fluorescence autocorrelation curves to a model that takes into account both the translational diffusion process of either a single or two diffusional components and a rapid fluorescence fluctuation process. The fitting procedure accounted for the bulk viscosity at different PEG concentrations, as well as the effective excitation volume, which was estimated using free ATTO 532 dye (Figure S24; see Materials and Methods). These results are presented in Figure 7.

**Figure 7:**
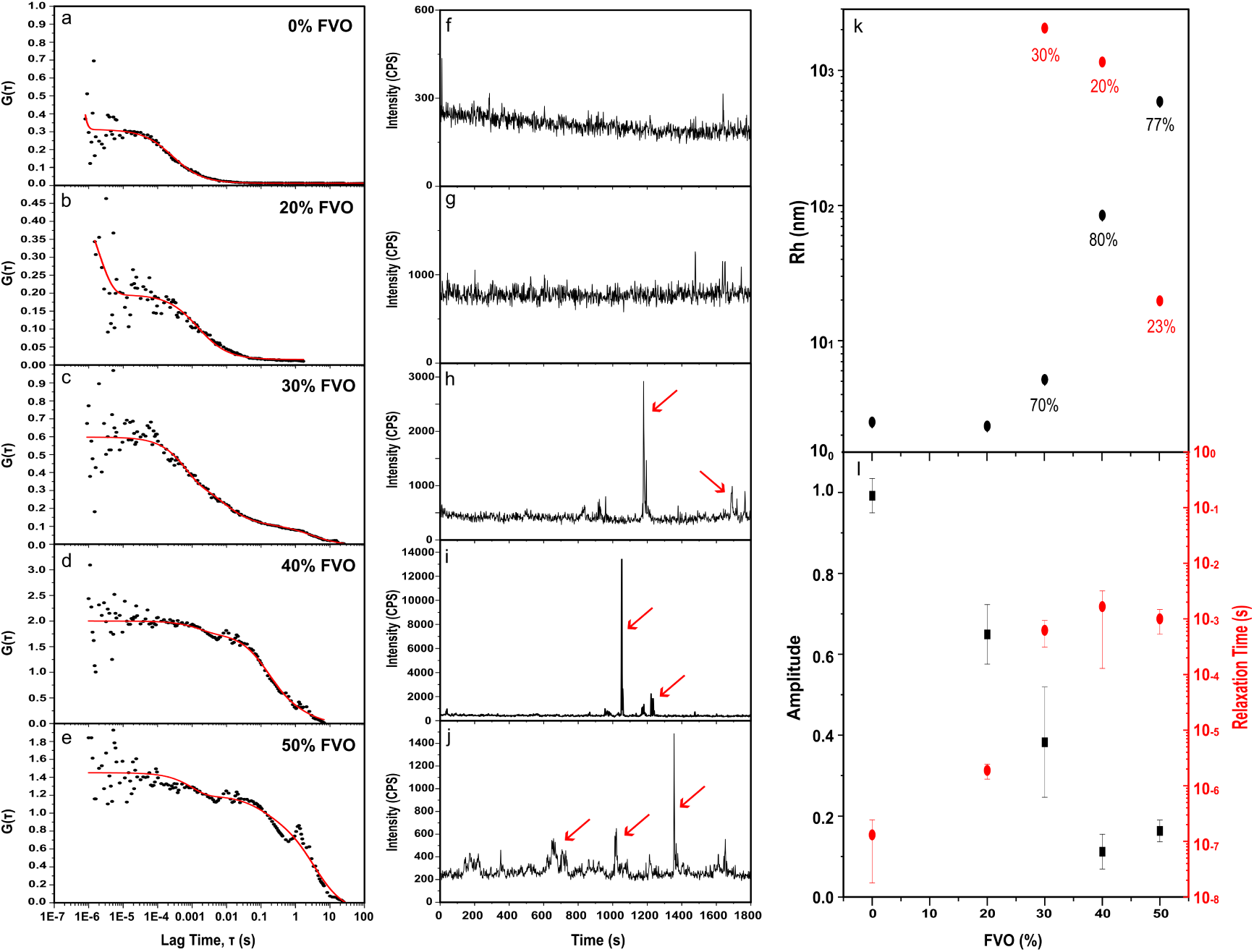
FCS measurements of mCherry fluorescence dynamic processes. The fluorescence autocorrelation curves (a-e, black) of fluorescence traces of 1 nM mCherry (f-j, black) in PBS buffer with 0% (a, f), 20% (b, g), 30% (c, h), 40% (d, i) & 50% (e, j) FVO of PEG 6,000, together with the best-fit theoretical curves (a-e, red). Hydrodynamic radius values are calculated from the best-fit diffusion coefficient values (k) using the Stokes-Einstein equation (Eq. 5), after factoring out the viscosity of PEG. The hydrodynamic radii are shown for each % FVO PEG for fits to FCS models of single (solely black) or two (black and red) diffusion components (Eq. 3). In fits to a model of two diffusion components, the fraction of the diffusion component is provided in the figure panel. The best-fit parameter values are also shown for the rapid fluorescence fluctuation process (l). Red arrows point at high fluorescence burst events corresponding to mCherry aggregates.

Using the best-fit values of the diffusion coefficients from FCS model fittings (Figure 7 a-e) and the Stokes-Einstein equation (Equation 5), the hydrodynamic radii were calculated for each PEG concentration either for a single diffusion component model (Figure 7k, black) or for a two diffusion components model (Figure 7k, black and red). The hydrodynamic radius of mCherry calculated from the best-fit values of the diffusion coefficient to the FCS results, in the absence of PEG and in the presence of up to 20% FVO of PEG, is ∼2.33-2.50 nm. Interestingly, above 30% FVO of PEG, there is an increase in the mean hydrodynamic radius from twofold up to ∼2 orders of magnitude, corresponding to the appearance of mCherry aggregates. We also observe the appearance of high fluorescence bursts in the fluorescence traces at elevated PEG concentrations (see Figure 7h-j, red arrows). Since mCherry in its folded state is very stable ^112^ (Figure S25), it is highly unlikely that the aggregates are composed of misfolded mCherry molecules. Moreover, if they were misfolded proteins that aggregated, the environment of their chromophore would deform, leading to the loss of fluorescence, which is not the case here. Alternatively, the PEG-mCherry interaction identified in the simulations (Figure 5) could support bringing multiple mCherry proteins together to form large aggregates. In particular, PEG-wrapping around the side chains of surface-exposed lysine residues could support the aggregation of mCherry. Notably, the PEG concentrations for these aggregates to emerge occur at 30% FVO of PEG, which is the same threshold previously found for fluorescence lifetime changes of mCherry ^50^.

Based on the best-fit values of the rapid fluorescence fluctuation process (Figure 7l), we can speculate whether trends in the timescale and efficiencies of the rapid fluorescence fluctuations as a function of PEG concentration might be related to the ns structural dynamics found in the simulations. Interestingly, we identify fluorescence fluctuations in the absence of PEG occurring in the hundreds of ns (Figure 7l, red), with a non-negligible equilibrium fraction (Figure 7l, black). This could mean that the reported *β*7-*β*10 gap dynamics might influence fluorescence fluctuations to some degree. However, at ≥ 30 % FVO of PEG, this rapid process slows down by 3-4 orders of magnitude (Figure 7l, red), and its equilibrium fraction decreases dramatically (Figure 7l, black). Here, we show that the chromophore can still have slower relaxation dynamics even when the average fluorescence lifetime decreases. The fluorescence lifetime decrease results from of an increase in the nonradiative de-excitation events ^50^. However, part of the overall relaxation time is the vibrational relaxation processes. While the average fluorescence lifetime decreases, the time it takes for the chromophore to relax to the lowest vibrational state of the ground state may be longer or even increase due to interactions between PEG and mCherry or as a result of the increased pocket volume, which leads to an increase in the degrees of freedom of the chromophore. These results qualitatively suggest that, in addition to the slowdown in structural dynamics observed in the simulations in the presence of elevated PEG concentrations (Figure 3), these structural dynamics do not affect the observed fluorescence fluctuations without PEG or at lower PEG concentrations. Although we have no way to identify the direct cause of the fluorescence fluctuations, it is likely to be the combined effect of multiple structural fluctuations in the vicinity of the chromophore, where the *β*7-*β*10 gap dynamics is one of the contributors. In the fluorescence data, the equilibrium fraction decreases dramatically above 30% PEG FVO, but this is not the equilibrium fraction of the *β*7-*β*10 gap states – it is the equilibrium fraction of a dim state of the fluorescence fluctuation process, and we do not know how the dim state relates to the *β*7-*β*10 gap states. Since we expect the presence of PEG to cause a slowdown of structural dynamics generally through its viscosity, this slowdown in fluorescence fluctuations is consistent with the slowed structural fluctuations observed in our simulations.

Altogether, the reported FCS results add corroboration to the findings about the PEG-mCherry interactions identified in the MD simulations and offer additional insights into the immediate implications of these interactions. The FCS results also suggest a potential connection between the observed ns structural dynamics and its potential influence on the fluorescence fluctuations arising from the chromophore itself. Both the FCS results and the results of the simulations are consistent with a slowdown of structural dynamics in the presence of PEG.

### Crowder Bulkiness and Chemical Structure, and Their Effects On mCherry Fluorescence Lifetime

Previously, we have shown that PEG concentrations above 30% FVO induce a reduction in mCherry fluorescence lifetime ^50^. While PEG is commonly used in *in vitro* studies as a crowding agent, other polymers are also commonly used, such as dextran or Ficoll, which are bulky, unlike the more flexible PEG. Here, we compare the changes in the mean fluorescence lifetime of mCherry as a function of % FVO of EG to the mean fluorescence lifetimes of mCherry as a function of the bulkier crowder, dextran, of similar sizes (Figure 8). While PEG is a linear polymer constructed of repeating subunits of ethylene glycol, dextran polymers, in contrast, are branched polysaccharides that include repeating subunits (Figure 8c). The bulkiness of the polymeric subunit may influence the degree to which the crowder interacts with the macromolecule experiencing crowding. Solutions including bulky crowders are expected to exhibit increased excluded volume compared to less bulky crowders. Indeed, in the presence of PEG, we find more and larger aggregates compared to solutions that include dextran (Figure S26). Therefore, we tested whether the bulkier dextran crowding agent, of a size similar to the PEG size used in our prior work, can induce mCherry fluorescence lifetime reduction. Our results show that, unlike with PEG, the mean fluorescence lifetime of mCherry does not monotonically reduce as a function of increasing dextran concentration (Figure 8a,b). This result exemplifies the importance of the softness of the crowder in crowder-mCherry interactions and suggests that soft interactions such as those seen here in MD simulations are an important factor in the observed decrease in mCherry’s fluorescence lifetime. While dextran, much like PEG, includes internal and terminal oxygen groups, the bulkiness of these groups may influence their surface exposure and, hence, the probability they will interact with surface-exposed protein residues.

**Figure 8:**
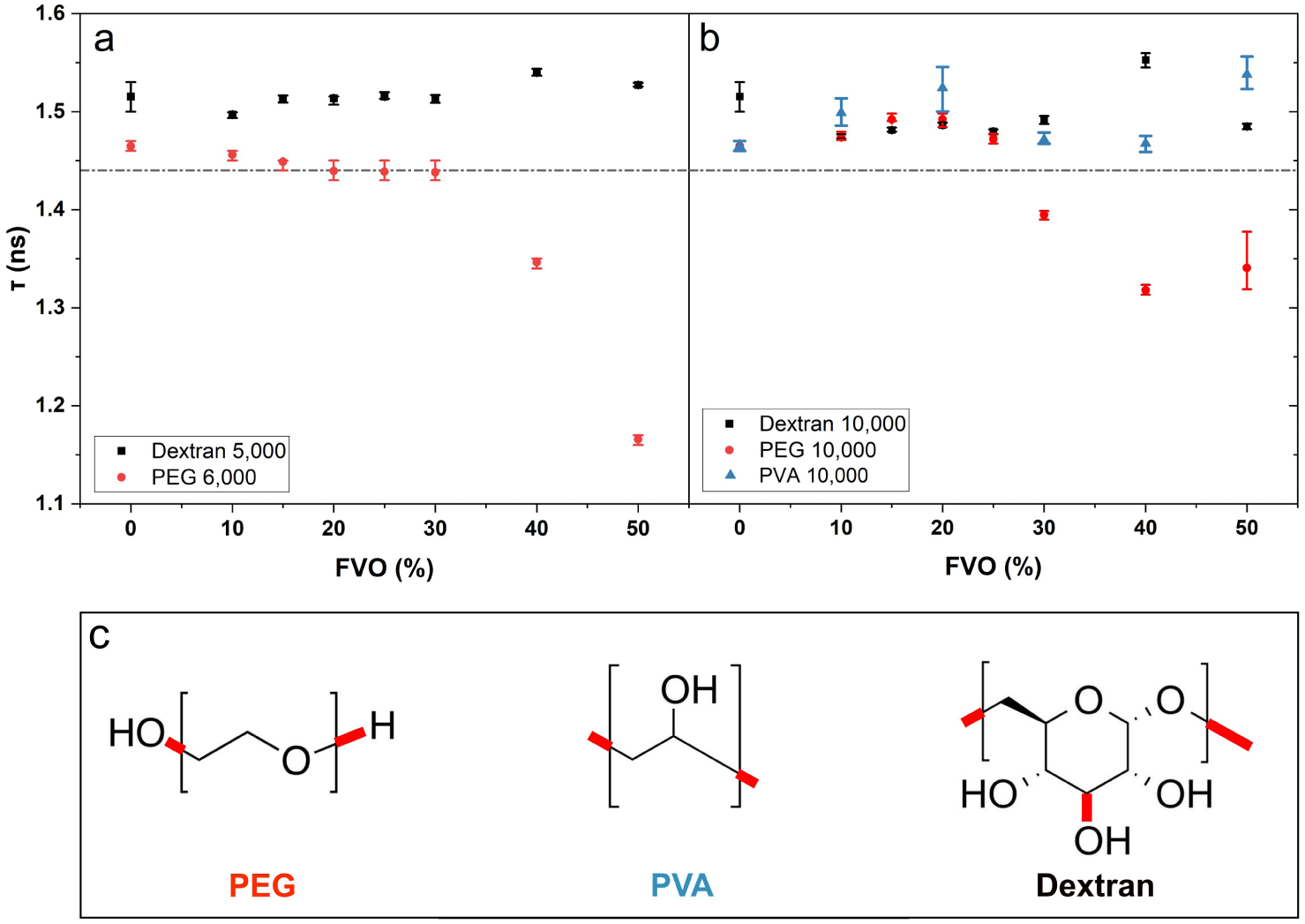
Mean fluorescence lifetime of mCherry in the presence of increasing % FVO of PEG, PVA, or dextran. The mean fluorescence lifetime (*τ* ; see Eq. 7) of 100 nM mCherry as a function of increasing (a) % FVO of dextran 5,000 (black) or PEG 6,000 (red), and (b) dextran 10,000 (black), PEG 10,000 (red), or PVA 10,000 (blue). The dashed line marks the lowest estimate of the mean fluorescence lifetime for mCherry without the influence of crowders ^50^. The chemical structures of a single subunit of each crowder are shown (c), and the polymer linking bonds are highlighted (c, red). Two linking bonds represent a linear polymer (e.g., PEG and PVA), and three linking bonds represent a branched polymer (e.g., dextran).

Importantly, the MD simulations suggest PEG has specific interactions with exposed positively-charged side chains in mCherry that could be stabilized through the chemical groups of the repeating subunits in PEG. To test this suggestion experimentally, we compared the influence of PEG on the mean fluorescence lifetimes of mCherry to those influenced by another linear polymer with subunits having a different chemical structure, polyvinyl alcohol (PVA; Figure 8b,c). Interestingly, unlike PEG, and much like the bulkier dextran, PVA does not induce the mean fluorescence lifetime reduction effect. This result strengthens the suggestion that it is not only the flexibility of a linear polymer but also the specific chemical structure of its subunits that contribute to the PEG-mCherry interactions that induce the fluorescence lifetime reduction.

To further investigate the effects of dextran crowding on mCherry, we performed FCS measurements with increasing concentrations of dextran (Figure S27). Similar to the results with PEG, we find that the hydrodynamic radius of mCherry in the presence of up to 20% FVO of dextran is ∼2.50 nm. Much like with PEG, we observe aggregates above 30% FVO dextran. These aggregates are smaller compared to aggregates that form in the presence of PEG of similar size and concentration (Figure S26). This could mean that while dextran induces crowding, similarly to PEG, the resulting aggregates do not become as large because interactions between mCherry proteins might not be mediated by interactions with the crowder, as in PEG-induced aggregates. Macromolecular crowding is known to enhance self-association solely through its excluded volume effects ^113^. However, additional crowder-crowdee interactions may mediate further enhancement of aggregation. Furthermore, unlike with PEG, we find that the timescale of rapid fluorescence fluctuations does not exhibit a clear monotonic change with increasing dextran concentration, but rather remains at hundreds of ns (Figure S27). Nevertheless, the equilibrium fraction of the rapid fluorescence fluctuations does seem to be influenced by the increase in dextran. This suggests that unlike with PEG, elevated dextran concentration does not affect the timescale of fluorescence fluctuations, potentially since it does not interact specifically with mCherry. Altogether, these results further support the notion that in addition to crowding, which is induced by PEG as well as dextran, specific PEG-mCherry interactions could mediate bringing multiple mCherry proteins together into forming larger and more stable aggregates, as well as influence the structural dynamics of mCherry. Elevated dextran concentrations also induce the formation of PEG aggregates. However, the fluorescence lifetime effect that occurs in the presence of PEG does not occur in the presence of dextran. Together, this indicates that mCherry aggregation is not the cause of the fluorescence lifetime effect seen in PEG. Additionally, and as has been noted by us before ^50^, although nonradiative excitation energy transfer between mCherry subunits, also known as homoFRET, can occur in the aggregate, homoFRET is not expressed as fluorescence lifetime changes. Therefore, it is unlikely that the fluorescence lifetime reduction observed in the presence of PEG occurs due to its aggregation.

## Discussion

This work provides a detailed description of the influence of PEG-induced crowding on the structure and dynamics of mCherry and provides insight into the nature of PEG-protein interactions. Since the effect of PEG on mCherry occurs on a structural level, we required an all-atom model with explicit solvent to probe changes in the structural dynamics of mCherry. Our simulations show that PEG wraps around solvent-exposed positively charged residues, consistent with previous studies ^102–106^. This binding mode, which appears to occur between positively charged NH_3_ groups that are distant from other positively charged residues, does not fit into the category of weak non-specific interactions and suggests that there may be more to PEG-protein interaction than excluded volume effects. Compared to other crowders, like dextran, which is bulky and stiff, the malleability of PEG facilitates its conformational rearrangement to enhance attractive interactions with proteins. However, one should take into account the chemical nature of the flexible linear polymer and not only the level of bulkiness when examining crowder-protein specific interactions such as those suggested with PEG and mCherry. The comparison of the effects of PEG versus PVA on the fluorescence lifetime changes in mCherry suggests that, indeed, PEG-mCherry interactions and their influence on the fluorescence characteristics of mCherry are specific. The tendency of PEG to favour interactions with specific protein residues more than others indicates that it cannot be described as a classical rigid body excluded volume crowder nor as a simple viscogen.

Consistent with previous work, we observe large structural changes in the short disordered regions of mCherry and subtle changes in the global structure of its ordered regions. PEG induces increased helicity and compaction of the disordered regions of mCherry, in line with the trends seen for other IDRs, both short and long ones. Additionally, we observe local expansion of certain parts of the structured region of the protein in the presence of PEG. In the context of volume exclusion and soft interactions, on one hand, we suppose that the volume exclusion effect would have little influence on the structure of the *β*-barrel, since such a highly ordered and thermodynamically stable structure has little access to extended states even in the absence of crowders. On the other hand, we observe clear evidence of attractive interactions between PEG and mCherry that exert an overall outward force on the barrel and could explain the subtle overall expansion of the chromophore pocket. We speculate that a similar mechanism could be present for other well-folded proteins that have a predominant compact conformation, where the excluded volume effect is negligible compared to the influence of attractive soft interactions. In summary, the physical and chemical nature of the crowder molecule at high concentrations is important in directly affecting the structure and interactions of proteins rather than just influencing their conformational equilibria via excluded volume effects.

Regarding the observed aggregation and precipitation of mCherry in the presence of PEG, it is well known that PEG precipitates proteins ^114;115^. In addition, PEG has been widely used in studies testing the propensity for liquid-liquid phase separation (LLPS) of minimal protein and RNA components *in vitro*. To observe LLPS-like behaviour in these studies, increased concentrations of biomolecules are tested, with PEG included as a crowding agent. Biologically relevant LLPS is supported by weak multivalent interactions between IDRs above a given saturating concentration ^116–118^. Many proteins with IDRs have been shown to undergo PEG-induced LLPS, with *α*-synuclein being a representative example ^119^ with important biological consequences of its self-associated species. While this study has also shown similar behaviour in cells, it does not yet mean that the solution-based PEG-induced species fully represent the in-cell findings. In a recent study, the solution-based LLPS behaviour of *α*-synuclein was tested when the protein was tagged with either a small organic dye or with different FPs (including mCherry). Differences in LLPS-like behaviour were observed in the presence of PEG depending on the fluorescent tag used ^120^. Importantly, in this work and in our previous study ^50^, we observed that mCherry, a well-folded and highly thermodynamically stable protein, exhibits solution-based LLPS-like behaviour in the presence of PEG. The interactions of PEG with mCherry uncovered here could support the formation of precipitates without the long IDRs typically required to form weak, multivalent interactions in biologically-relevant LLPS. On the basis of these observations, we suggest that solution-based LLPS behaviour should be studied carefully, and if possible, without inducing it with PEG, but only with elevated concentrations of the IDRs of interest in solution.

## Author Contributions

**Liam Haas-Neill**: Conceptualization, Formal Analysis, Investigation, Methodology, Software, Visualization, Writing Original Draft Preparation; **Khalil Joron**: Conceptualization, Formal Analysis, Investigation, Methodology, Visualization, Writing - Original Draft Preparation; **Eitan Lerner**: Conceptualization, Formal Analysis, Funding Acquisition, Project Administration, Supervision, Writing - Review & Editing; **Sarah Rauscher**: Conceptualization, Formal Analysis, Funding Acquisition, Project Administration, Supervision, Writing - Review & Editing

## Acknowledgments

This research was supported by a Connaught New Researcher Award to S.R., a Natural Science and Engineering Research Council of Canada (NSERC) Discovery Grant to S.R., and the Israel Science Foundation to E.L. (grants 556/22 and 3565/20). This research was enabled in part by support provided by Calcul Quebec (www.calculquebec.ca) and the Digital Research Alliance of Canada (alliance.can.ca).

## Data Availability

MD simulation data (initial coordinates and simulation trajectories) are provided for all systems as a Zenodo repository accessible here: https://doi.org/10.5281/zenodo.13904991. Data underlying figures are also available in this Zenodo repository. We also provide a list of all PDB IDs that contain PEG interacting with lysine residues.

## Description of Additional Files

### Video 1

Dynamics of the interaction of PEG molecules with the protein mCherry are shown. mCherry is shown in a surface representation. Positively charged residues are shown in blue, negatively charged residues are shown in red, polar residues are shown in green, and non-polar residues are shown in grey. PEG molecules are shown in van der Waals representation with carbon, oxygen, and hydrogen atoms shown in cyan, red, and white, respectively. The movie shows 2 *µ*s of simulation, and 1 second corresponds to 60 ns of simulated time.

## Notes

https://doi.org/10.5281/zenodo.10798189

### Competing Interest Statement

The authors have declared no competing interest.

### Summary of Updates

Minor revisions have been made throughout the main text and supporting information.

