## Supporting Information File for "PEG-mCherry interactions beyond classical macromolecular crowding"

### Contents

### List of Figures

S1 Sequence of mCherry construct. . . . . . . . . . . . . . . . . . . . . . . . . . . . . . . S4
S2 Secondary Structure of mCherry. . . . . . . . . . . . . . . . . . . . . . . . . . . . . . S6
S3 Time-series of mCherry structural features. . . . . . . . . . . . . . . . . . . . . . . . . S7
S4 Single exponential fits to autocorrelation. . . . . . . . . . . . . . . . . . . . . . . . . . S8
S5 Interactions with Periodic Images. . . . . . . . . . . . . . . . . . . . . . . . . . . . . S9
S6 Structural Distributions With and Without Periodic Image Interactions. . . . . . . . S10
S7 Distribution of $R_g$ for the structured region and tails of mCherry. . . . . . . . . . . . S11
S8 Difference map for inter-residue contacts. . . . . . . . . . . . . . . . . . . . . . . . . S12
S9 Distribution of Q163 $\chi_2$ and $\chi_3$ dihedral angles. . . . . . . . . . . . . . . . . . . . . . . S13
S10 Distribution of the pocket volume separated by open and closed state $\beta$7-$\beta$10 gap states.S14
S11 Joint distributions of structural properties and interactions with PEG. . . . . . . . . S15
S12 RMSF of N- and C- terminal IDRs. . . . . . . . . . . . . . . . . . . . . . . . . . . . S16
S13 Number of PEG molecules in contact with mCherry. . . . . . . . . . . . . . . . . . . S17
S14 PEG contact frequency by residue. . . . . . . . . . . . . . . . . . . . . . . . . . . . S18
S15 PEG contact frequency vs. solvent accessibility. . . . . . . . . . . . . . . . . . . . . S19
S16 Contacts with PEG by residue type. . . . . . . . . . . . . . . . . . . . . . . . . . . . S20
S17 Time series of K47-PEG interactions. . . . . . . . . . . . . . . . . . . . . . . . . . . S21
S18 PEG wrapping observed in mCherry crystal structure. . . . . . . . . . . . . . . . . . S22
S19 PEG wrapping varies strongly between lysine residues on the surface of mCherry. . . S23
S20 PEG wrapping in mCherry depends on local environment. . . . . . . . . . . . . . . . S24
S21 PEG wrapping in the anti-PEG antibody depends on the local environment. . . . . . S25
S22 Examples of PEG wrapping in the PDB. . . . . . . . . . . . . . . . . . . . . . . . . . S26
S23 Contacts between each monomeric unit of PEG and mCherry. . . . . . . . . . . . . . S27
S24 FCS best fit result for ATTO 532. . . . . . . . . . . . . . . . . . . . . . . . . . . . S28
S25 Thermal stability of mCherry. . . . . . . . . . . . . . . . . . . . . . . . . . . . . . . S29

S26 mCherry aggregates in PEG vs dextran. . . . . . . . . . . . . . . . . . . . . . . . . . S30
S27 FCS measurements of mCherry in increasing dextran concentrations. . . . . . . . . . S31

### List of Tables

S1 Simulation Systems . . . . . . . . . . . . . . . . . . . . . . . . . . . . . . . . . . . . . S4
S2 Chromophore pocket residues. . . . . . . . . . . . . . . . . . . . . . . . . . . . . . . . . S5
S3 mCherry autocorrelation times. . . . . . . . . . . . . . . . . . . . . . . . . . . . . . . . S5

| Trajectory | #Atoms | Box dimensions (nm) | #Water Molecules | PEG Concentration (% FVO) | NaCl Concentration (mM) |
|---|---|---|---|---|---|
| 1 | 38774 | 7.3 7.0 7.3 | 11632 | 0 | 139.9 |
| 2 | 38774 | 7.3 7.0 7.3 | 11632 | 0 | 139.9 |
| 3 | 38774 | 7.3 7.1 7.3 | 11632 | 0 | 139.9 |
| 4 | 38774 | 7.3 7.1 7.3 | 11632 | 0 | 139.9 |
| 5 | 38774 | 7.3 7.1 7.3 | 11632 | 0 | 139.9 |
| 6 | 36353 | 7.1 6.8 7.1 | 6713 | 42 | 139.9 |
| 7 | 36353 | 7.1 6.8 7.1 | 6713 | 42 | 139.9 |
| 8 | 36353 | 7.1 6.8 7.1 | 6713 | 42 | 139.9 |
| 9 | 36353 | 7.1 6.8 7.1 | 6713 | 42 | 139.9 |
| 10 | 36353 | 7.1 6.8 7.1 | 6713 | 42 | 139.9 |

**Table S1:** Simulations Systems. Each trajectory is 2 $\mu$s of simulation time for a total sampling time of 20 $\mu$s.

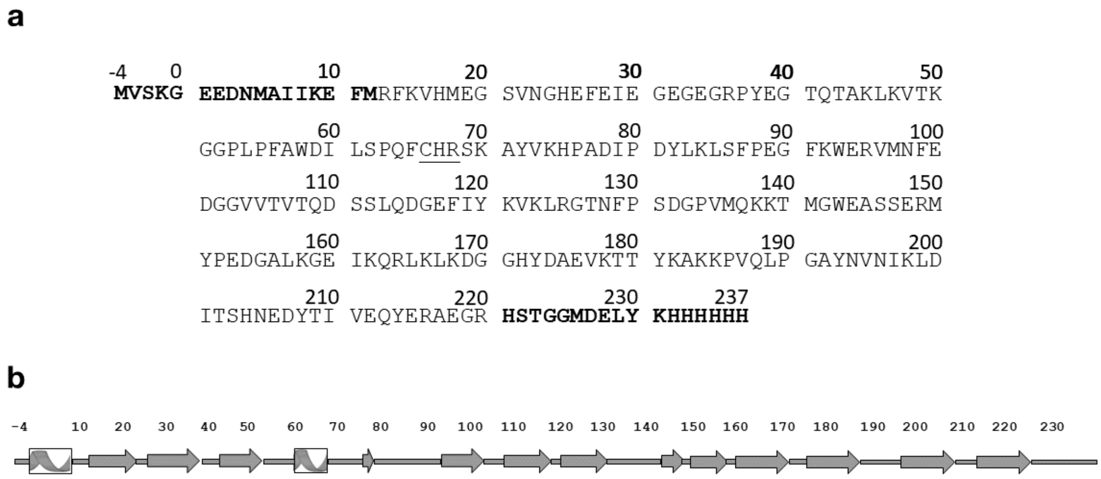


**Figure S1: Sequence of mCherry construct.** (a) The sequence of the mCherry construct used in simulations and experiments. Residue numbering begins from -4 to match the numbering in the PDB entry 2H5Q [1]. Residues that are part of the IDRs at the N- and C-termini are indicated in bold. The chromophore is indicated as CHR and is underlined. (b) The secondary structure from the simulations (Figure S2) with the residue numbers aligned.

| Residue | % of frames without PEG | % of frames with PEG |
|---|---|---|
| PHE14 | 84.7 | 77.6 |
| GLN42 | 99.9 | 99.5 |
| THR43 | 88.3 | 75.7 |
| ALA44 | 99.8 | 99.5 |
| LEU46 | 6.3 | 8.7 |
| SER62 | 16.8 | 24.6 |
| PRO63 | 99.9 | 100.0 |
| GLN64 | 87.2 | 84.9 |
| PHE65 | 100.0 | 100.0 |
| SER69 | 100.0 | 100.0 |
| LYS70 | 65.7 | 63.4 |
| PHE91 | 3.1 | 4.3 |
| TRP93 | 99.3 | 98.6 |
| ARG95 | 100.0 | 100.0 |
| GLN109 | 49.9 | 51.6 |
| TRP143 | 31.7 | 29.0 |
| GLU144 | 2.4 | 1.5 |
| ALA145 | 15.3 | 17.2 |
| SER146 | 41.1 | 47.8 |
| GLU148 | 4.2 | 7.1 |
| ILE161 | 95.4 | 99.4 |
| GLN163 | 85.7 | 83.0 |
| VAL177 | 0.7 | 1.4 |
| TYR181 | 29.0 | 37.4 |
| ILE197 | 99.5 | 99.6 |
| LYS198 | 3.7 | 1.7 |
| LEU199 | 100.0 | 100.0 |
| GLN213 | 99.7 | 99.8 |
| TYR214 | 57.1 | 49.2 |
| GLU215 | 100.0 | 100.0 |

**Table S2: Chromophore pocket residues.** The percentage of the simulation frames that the pocket residues are in contact with the chromophore is provided for each system. Residues that form contacts with the chromophore for less than 0.33 % of the simulation frames (VAL16, TYR120, ASP200, ARG216, HIS232, HIS233, HIS236) are not listed in the table and are not considered to be part of the chromophore pocket.

| System | Feature | $\tau_{slow}$ (ns) | $\tau_{fast}$ (ns) | $\tau_{avg}$ (ns) | Amplitude |
|---|---|---|---|---|---|
| no-PEG | $\beta$7-$\beta$10 Gap Width | 69.87±1.13 | 1.09±0.04 | 42.9±0.7 | 0.39 |
| PEG | $\beta$7-$\beta$10 Gap Width | 100.20±7.39 | 0.97±0.36 | 71.5±5.3 | 0.29 |
| no-PEG | Q163 $\chi$1 | 19.61±6.17 | 2.50±0.69 | 11.1±3.1 | 0.50 |
| PEG | Q163 $\chi$1 | 47.42±12.99 | 4.13±0.39 | 21.3±5.2 | 0.60 |

**Table S3:** Parameters of the sum of two exponentials fit to autocorrelations. The fits were modelled according to Equation (1). $\tau_{avg}$ was computed using Equation (2). Fits for both the Q163 side chain and the $\beta$7-$\beta$10 gap width are shown with and without PEG.

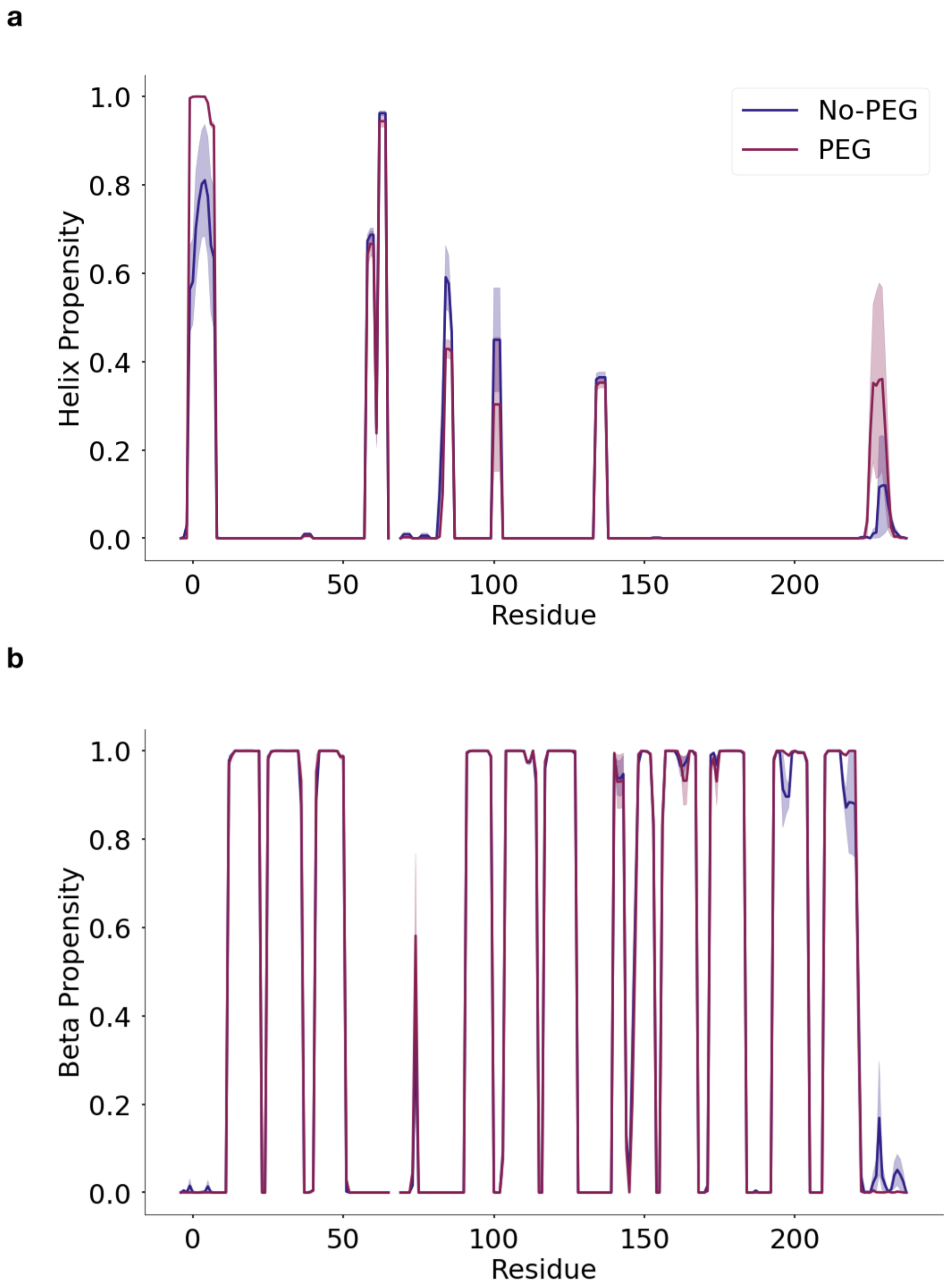


**Figure S2: Secondary Structure of mCherry.** The secondary structure of mCherry in the simulations trajectories is shown. The helix propensity (a) and beta propensity (b) by residue are shown for the simulations with (plum) and without PEG (navy). Secondary structure propensity was computed as the portion of frames that a given residue is in a particular secondary structure state according to the DSSP algorithm [2]. Shading indicates the standard error of the mean obtained by treating each of the five trajectories as independent.

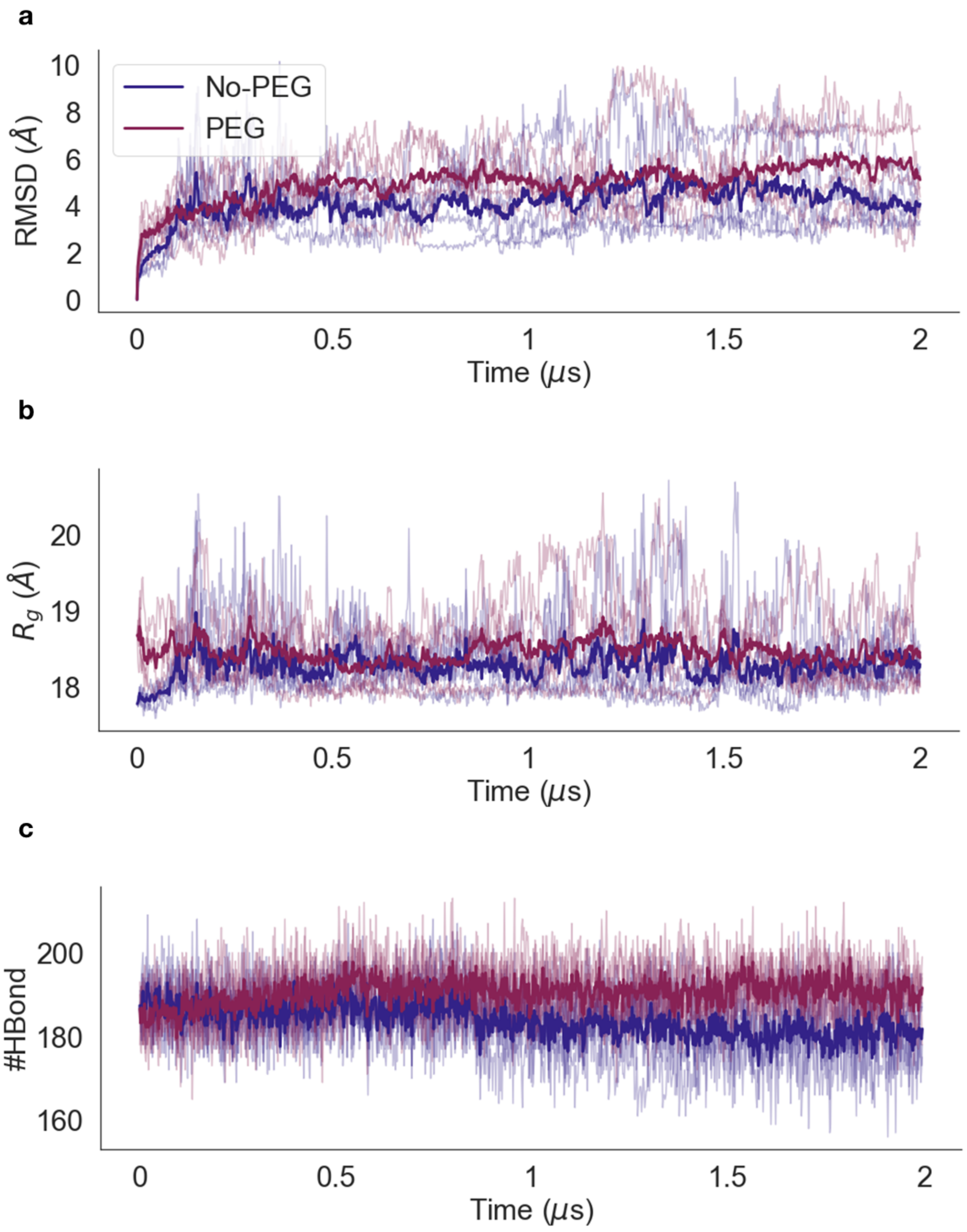


**Figure S3: Time-series of mCherry structural features.** (a) Time-series of the backbone RMSD for mCherry with and without PEG. (b) Time-series of the $R_g$ for mCherry with and without PEG. (c) Time-series of the number of intra-protein hydrogen bonds for mCherry with and without PEG. All trajectories are shown individually, the mean over trajectories for each system is shown with a darker line. The system with PEG is shown in plum and the system without PEG is shown in navy in each figure panel.

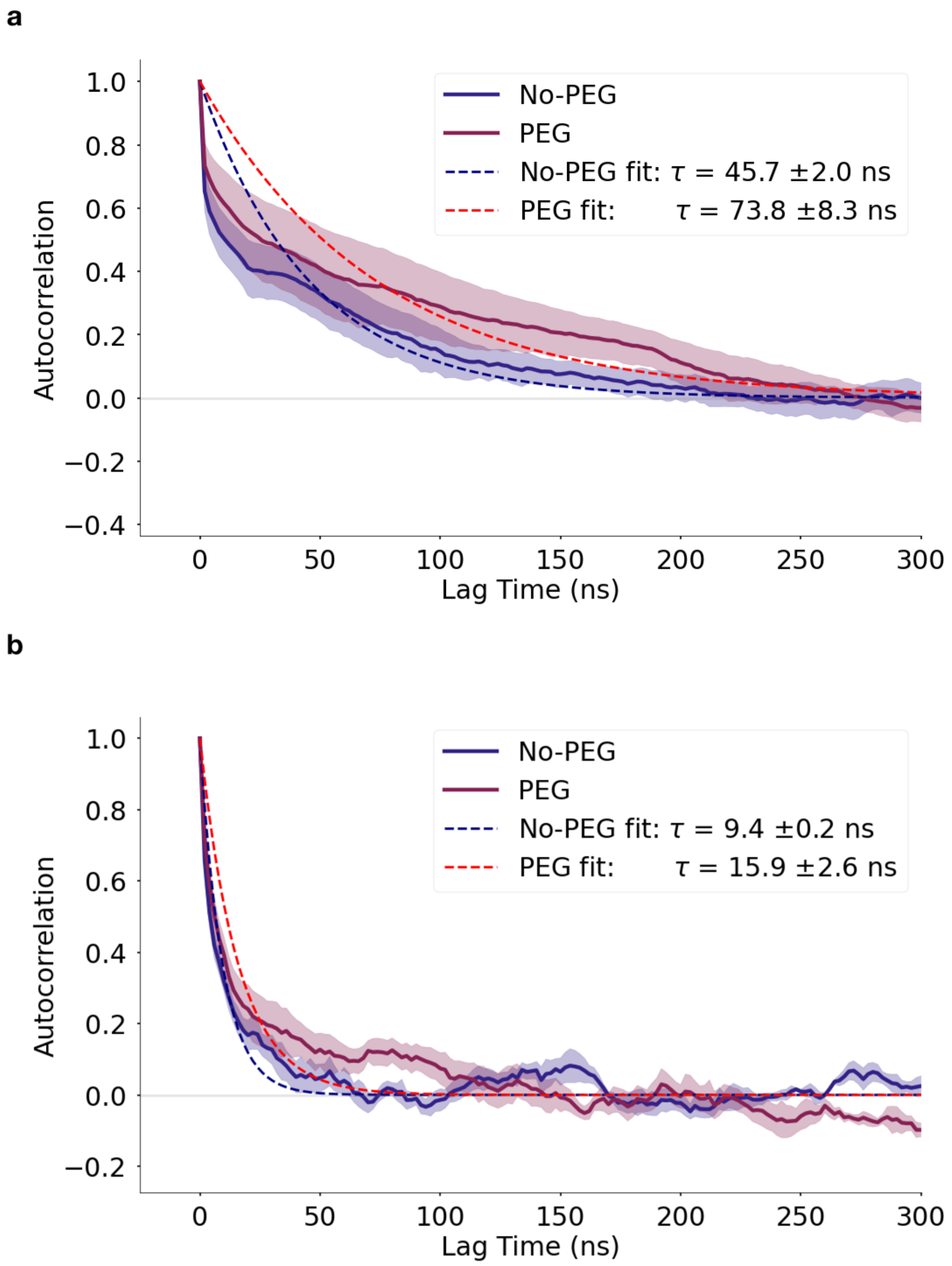


**Figure S4: Single exponential fits to autocorrelation.** The autocorrelation of the $\beta 7$-$\beta 10$ gap width (a) and Q163 dihedral angle (b) for both the PEG and no PEG systems. The single exponential fit to each autocorrelation is shown with the characteristic timescale, $\tau$, shown in the legend. The fit for the PEG system is shown in red, and the fit for the no-PEG system is shown in blue. The PEG system is shown in plum, and no-PEG is shown in blue. Shaded regions indicate the standard error of the mean, obtained by treating each trajectory as independent.

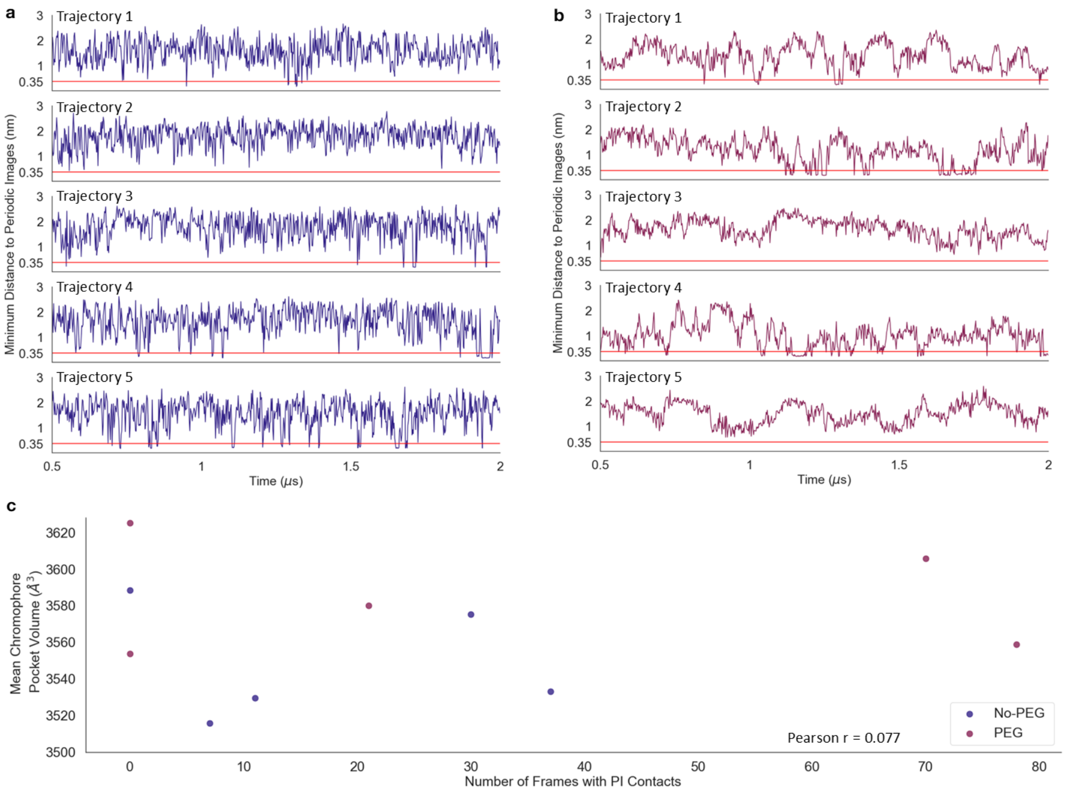


**Figure S5: Interactions with Periodic Images.** (a-b) Time series of the minimum distance between mCherry and its periodic images in each simulation trajectory. Trajectories without PEG are shown in navy (a) and trajectories with PEG are shown in plum (b). Red horizontal lines indicate 3.5 Å distance. (c) A scatter plot of the total number of frames in which contacts are formed between mCherry and its periodic images in a trajectory vs. the mean chromophore pocket volume. Each point represents a full trajectory. Trajectories with PEG are shown in plum and trajectories without PEG are shown in navy.

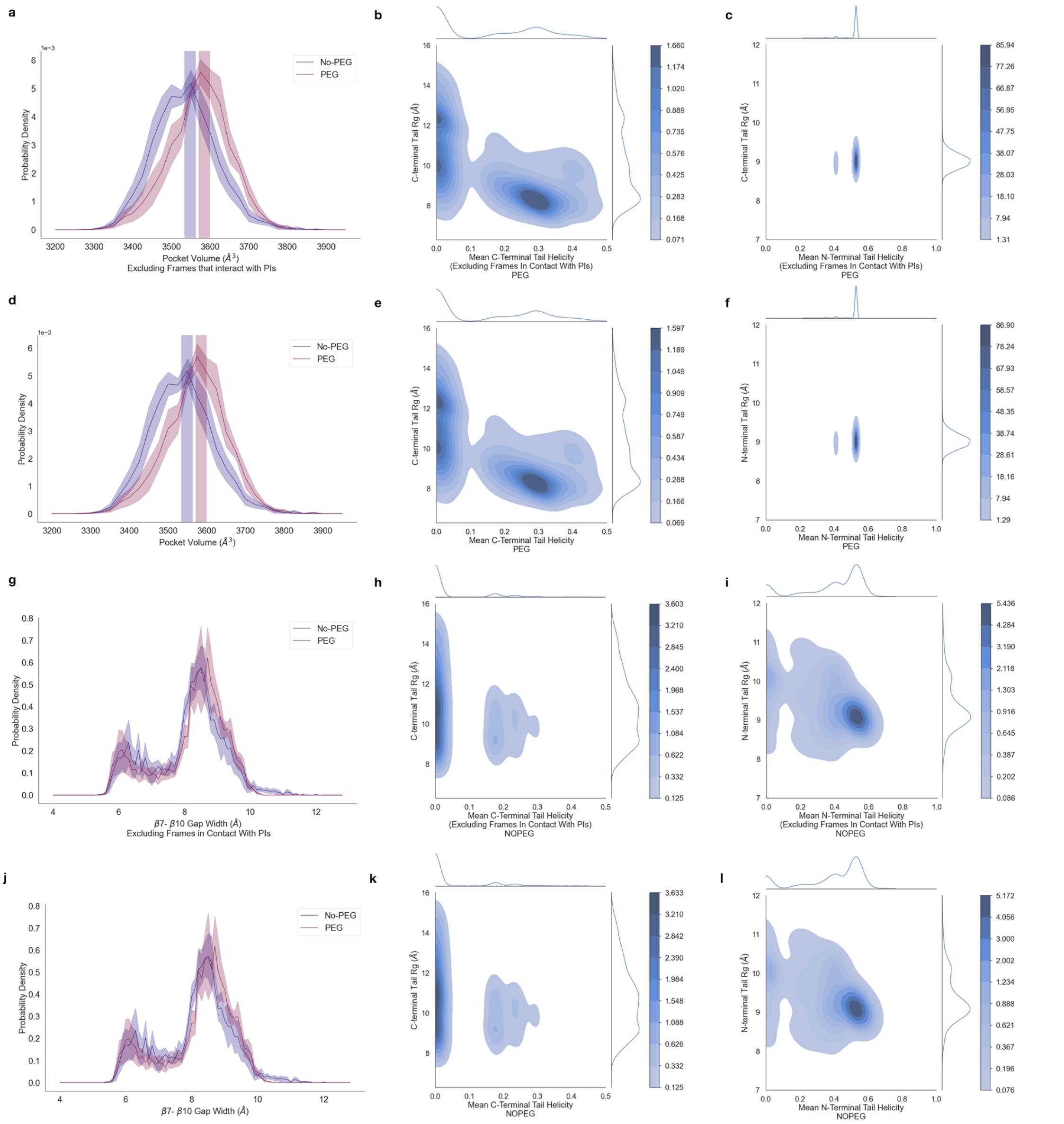


**Figure S6: Structural Distributions With and Without Periodic Image Interactions.** Distributions of various structural properties excluding frames that have contacts with periodic images (a-c, g-i) and including all frames (d-f, j-l). (a, d) Chromophore pocket volume. (b, e) C-terminal tail helicity and $R_g$ in PEG. (c, f) N-terminal tail helicity and $R_g$ in PEG. (g, j) $\beta$7-$\beta$10 gap width. (h, k) C-terminal tail helicity and $R_g$ without PEG. (i, l) N-terminal tail helicity and $R_g$ without PEG. In a, d, g, and j, distributions from PEG trajectories are shown in plum and distributions from trajectories without PEG are shown in navy. Shading indicates standard error of the mean.

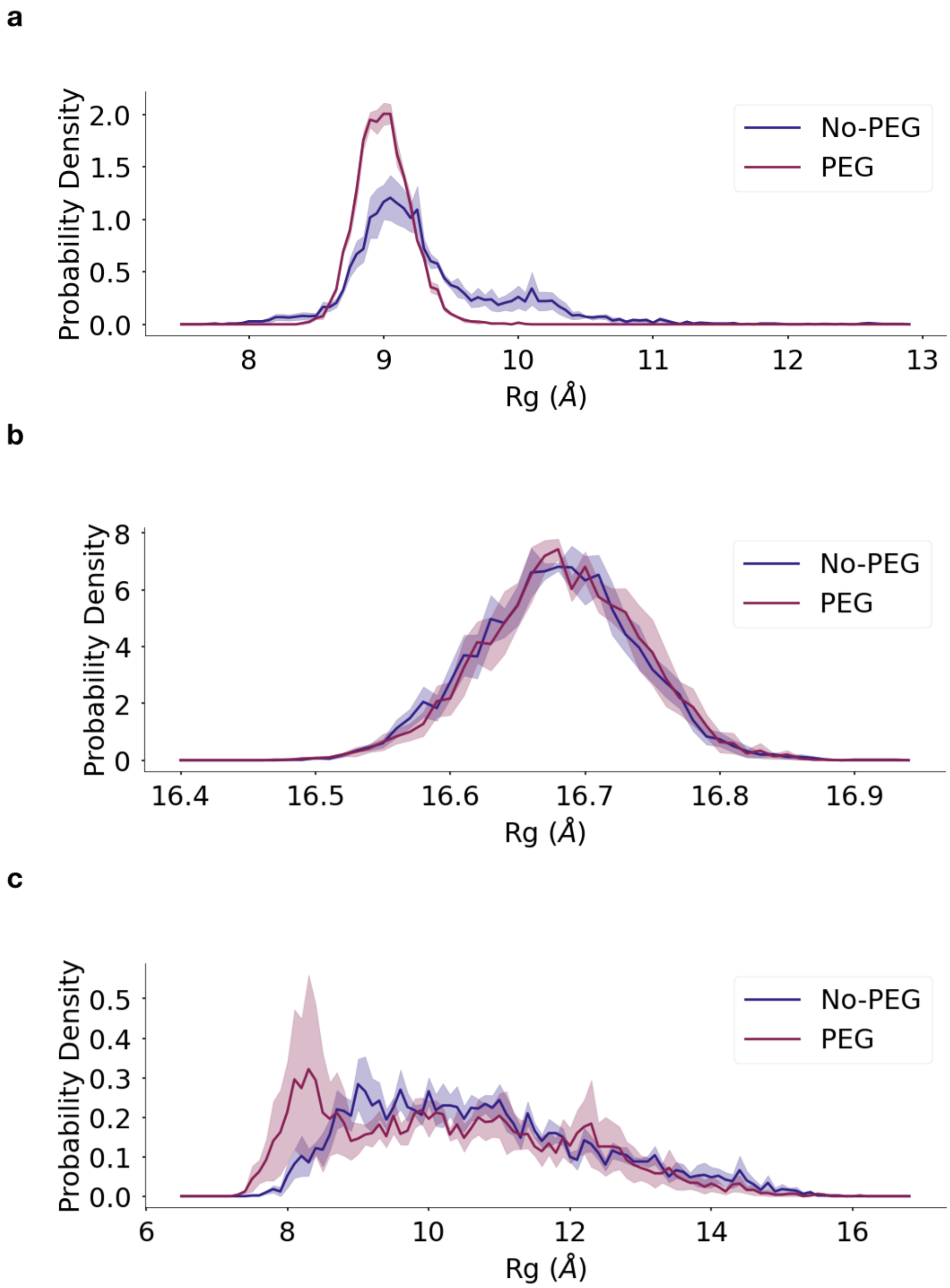


**Figure S7: Distribution of $R_g$ for the structured region and tails of mCherry.** (a) The distribution of $R_g$ for the N-terminal tail of mCherry (residues -4-12). (b) The distribution of $R_g$ for mCherry excluding the IDRs (residues 13-220). (c) The distribution of $R_g$ for the C-terminal tail of mCherry (residues 221-238). For all figure panels, the PEG system is shown in plum, and the no-PEG system is shown in blue. Shading indicates the standard error of the mean obtained by treating each of the five trajectories as independent.

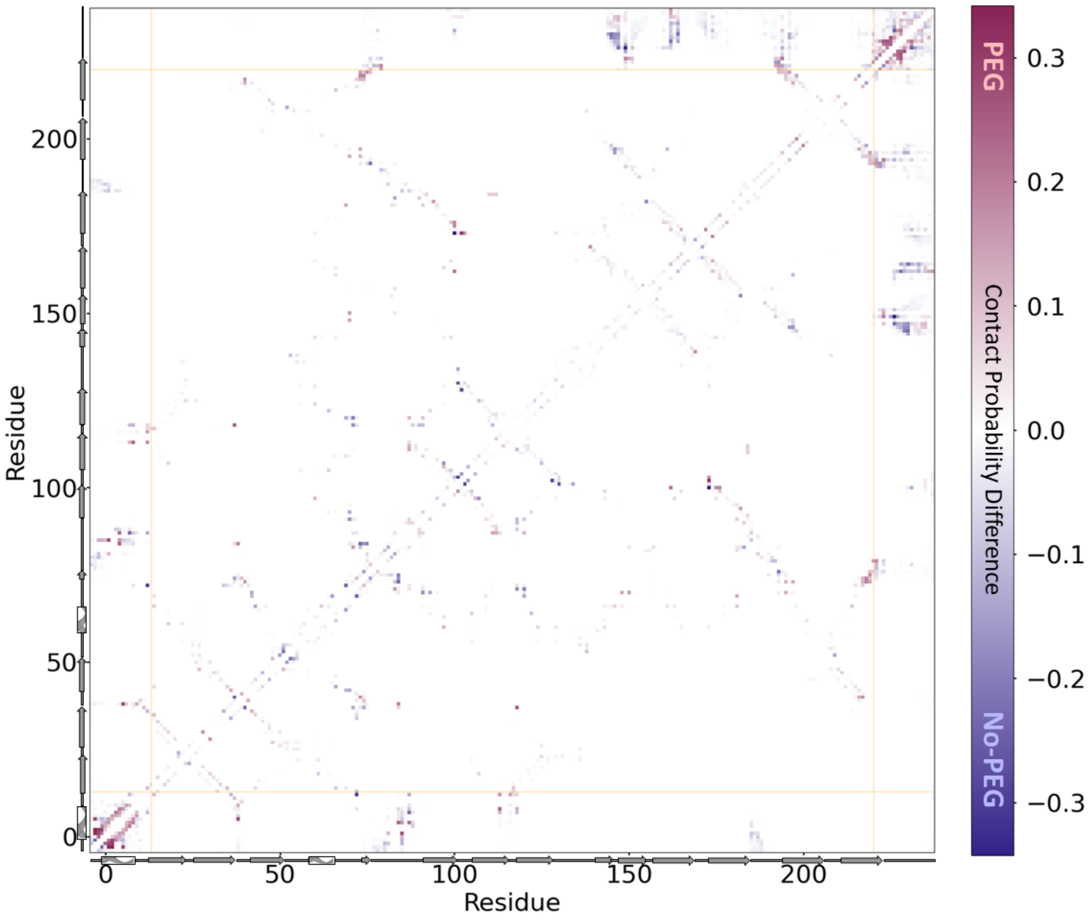


**Figure S8: Difference map for inter-residue contacts.** The contact difference map for the PEG system minus the no-PEG system. Contact probabilities were computed for each system and the difference was taken. Plum indicates that a contact is formed more often in PEG, while blue indicates that a contact is formed more often without PEG. The secondary structure is indicated on each axis. IDR boundaries are indicated by orange lines.

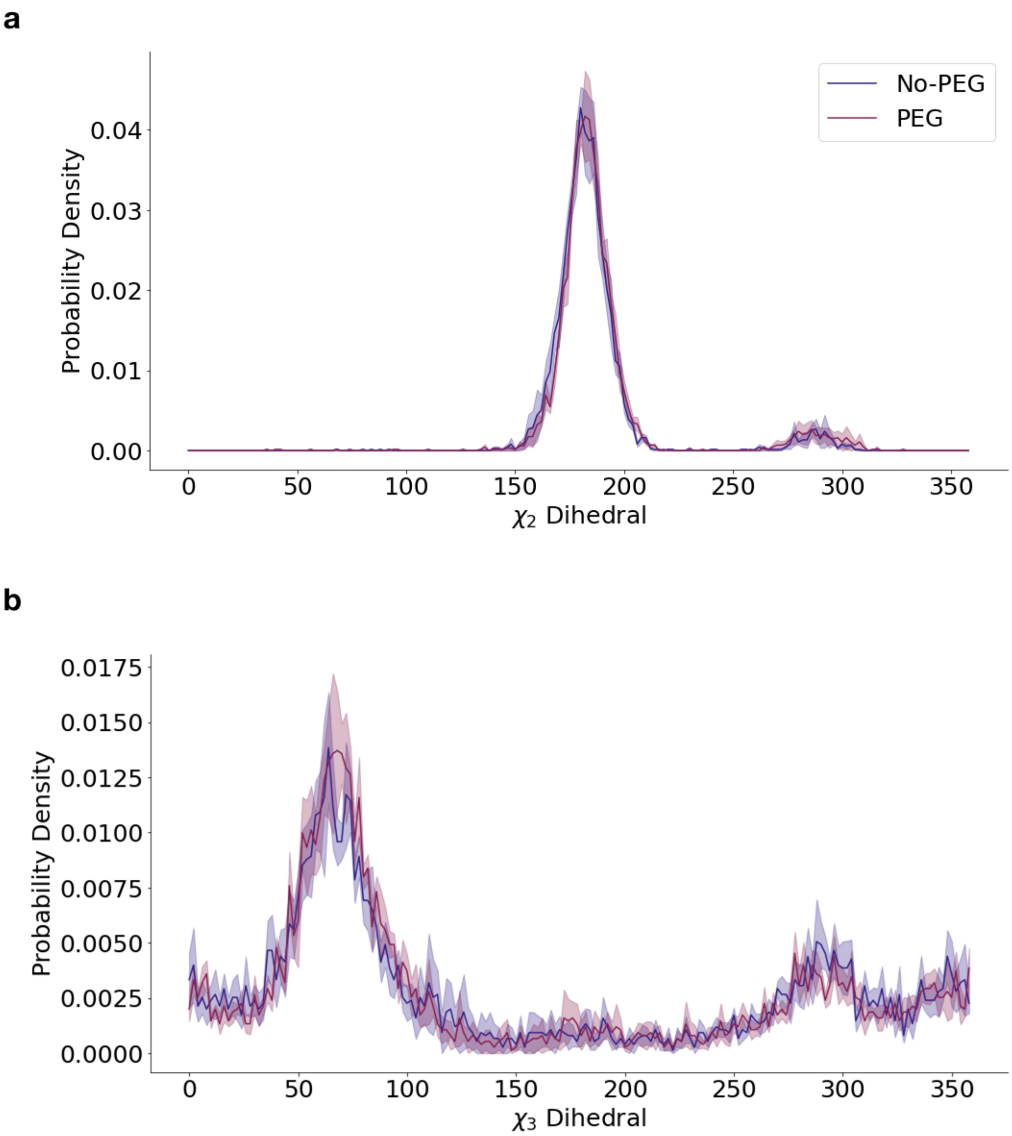


**Figure S9: Distribution of Q163 $\chi_2$ and $\chi_3$ dihedral angles.** (a) The distribution of Q163 $\chi_2$ dihedral angles with and without PEG. (b) The distribution of Q163 $\chi_3$ dihedral angles with and without PEG. The PEG system is shown in plum, and the no-PEG system is shown in blue. Shading indicates the standard error of the mean obtained by treating each of the five trajectories as independent.

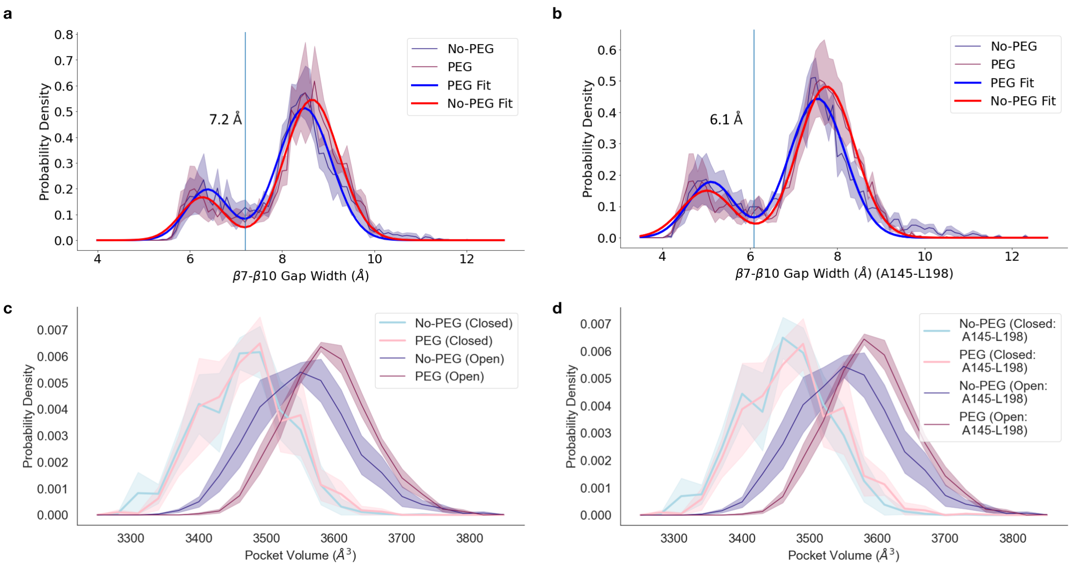


**Figure S10: Distribution of the pocket volume separated by open and closed state $\beta$7-$\beta$10 gap states.** (a-b) $\beta$7-$\beta$10 gap width distributions for the simulations with and without PEG using (a) our definition of the $\beta$7-$\beta$10 gap width and (b) using the Mukherjee *et. al.* definition of the $\beta$7-$\beta$10 gap width[3] The thick solid lines indicate the fit in each case (sum of two Gaussian functions), and the vertical blue lines indicate the local minimum in each fit that was used to define open and closed states of the $\beta$7-$\beta$10 gap. (c-d) The chromophore pocket volume distributions decomposed by $\beta$7-$\beta$10 gap state. States were assigned according to our definition of the $\beta$7-$\beta$10 gap width in (c) (open: $\beta$7-$\beta$10 gap width $\geq$ 7.2 Å, closed: $\beta$7-$\beta$10 gap width $\leq$ 7.2 Å) and according to the Mukherjee *et al.* definition[3] (open: $\beta$7-$\beta$10 gap width $\geq$ 6.1 Å, closed: $\beta$7-$\beta$10 gap width $\leq$ 6.1 Å) in (d). Distributions in the closed state are shown in lighter shades and distributions in the open states are shown in darker shades. Shading indicates the standard error of the mean obtained by treating each of the five trajectories as independent.

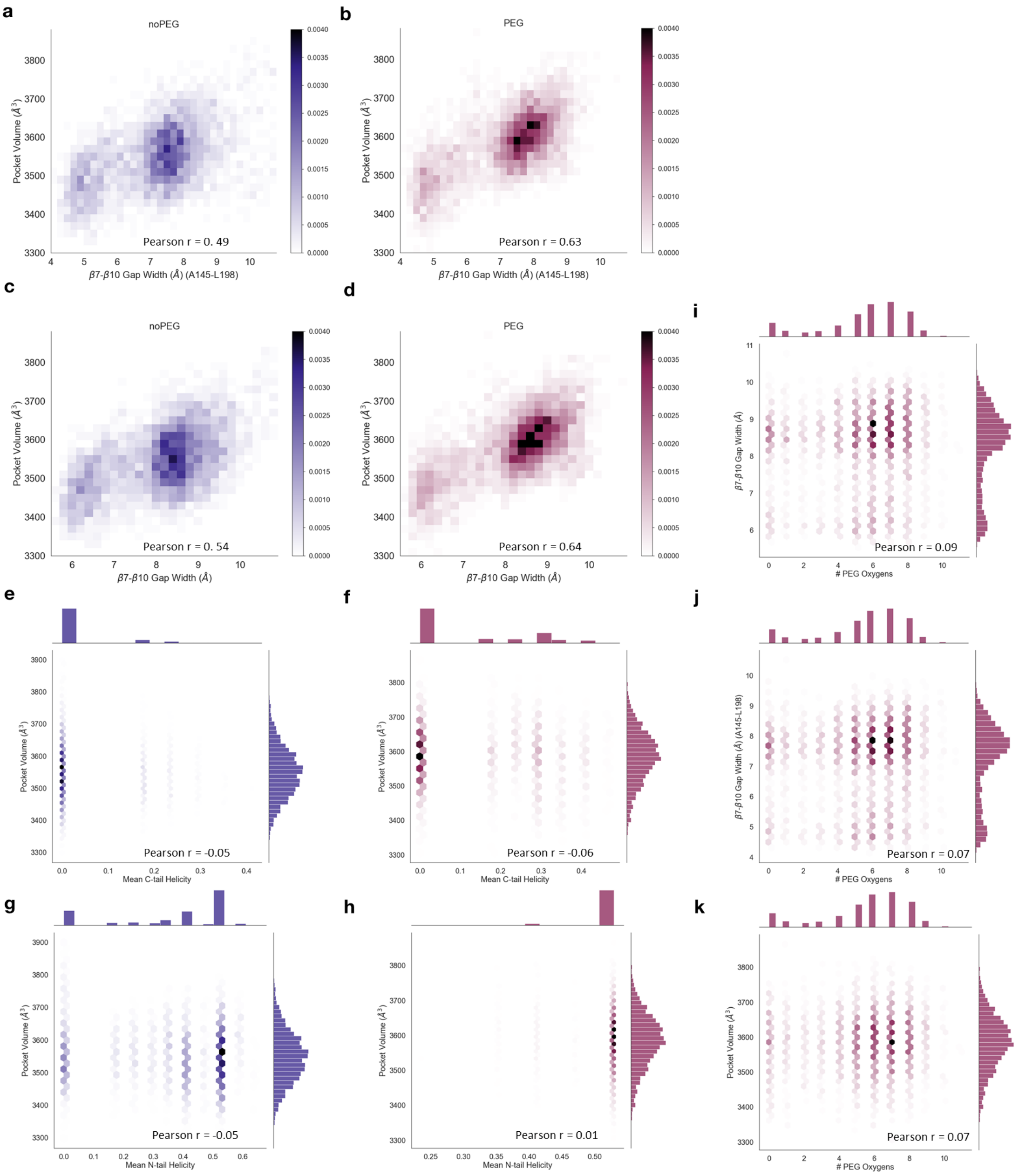


**Figure S11: Joint distributions of structural properties and interactions with PEG.** The joint distribution of the chromophore pocket volume and the width of the $\beta$7-$\beta$10 gap using the $\beta$7-$\beta$10 gap width definition of Mukherjee *et al.* for the simulations without PEG (a) and with PEG (b) and using our definition without PEG (c) and with PEG (d). The joint distribution of mean C-terminal tail helicity and the chromophore pocket volume for the simulations without PEG (e) and with PEG (f). The joint distribution of mean N-terminal tail helicity and the chromophore pocket volume for the simulations without PEG (g) and with PEG (h). The joint distribution of the number of PEG oxygens in contact with residue K47 and (i) the $\beta$7-$\beta$10 gap width using our definition (j) the $\beta$7-$\beta$10 gap width using the definition of Mukherjee *et al.*[3] (k) the chromophore pocket volume. Pearson r values for each pair of data sets are provided on each plot.

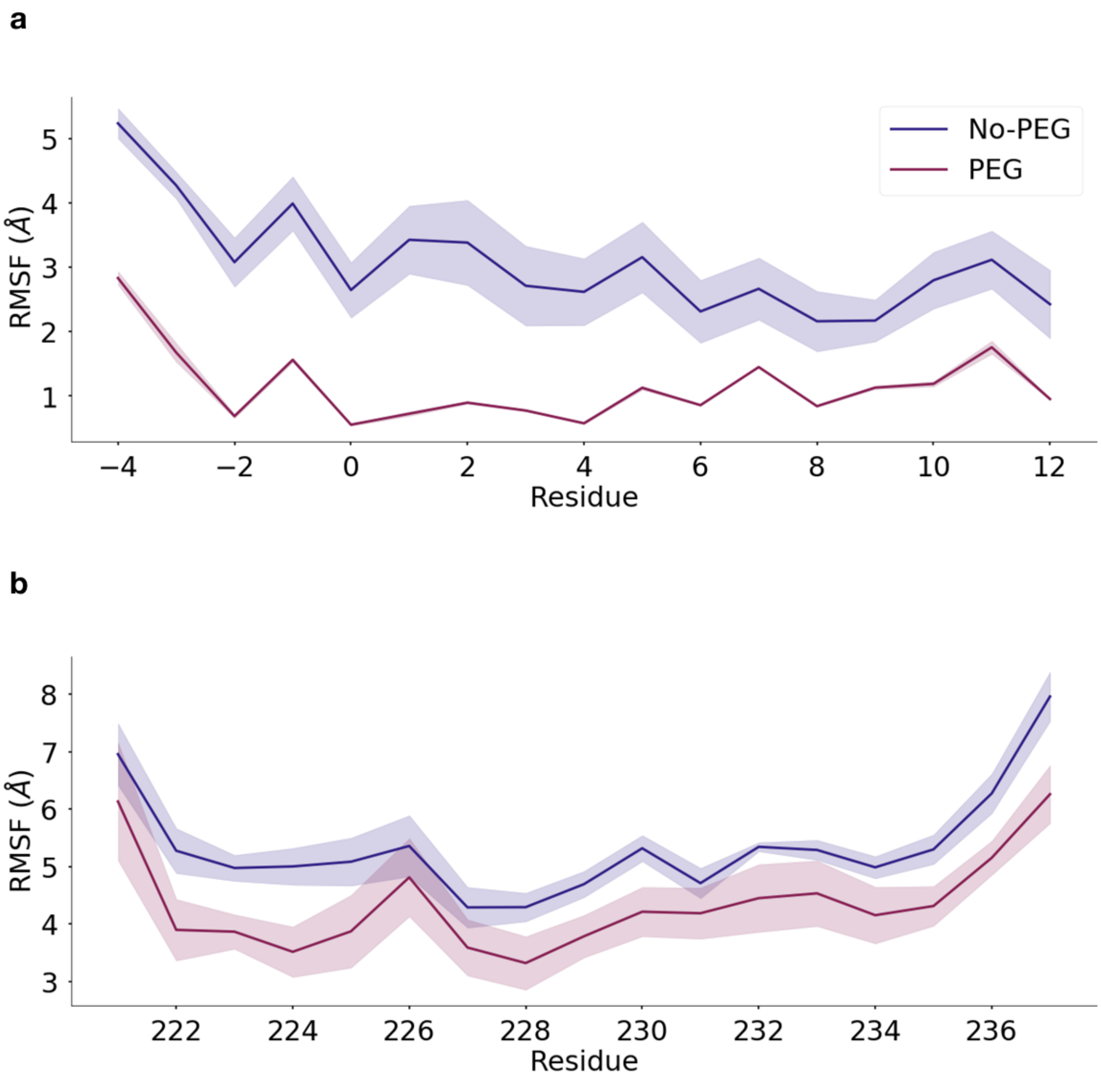


**Figure S12: RMSF of N- and C-terminal IDRs** (a) The RMSF including only the N-terminal tail is shown. (b) The RMSF including only the C-terminal tail is shown. The PEG system is shown in plum and the no-PEG system is shown in blue. Shading indicates standard error of the mean obtained by treating each trajectory as an independent measurement.

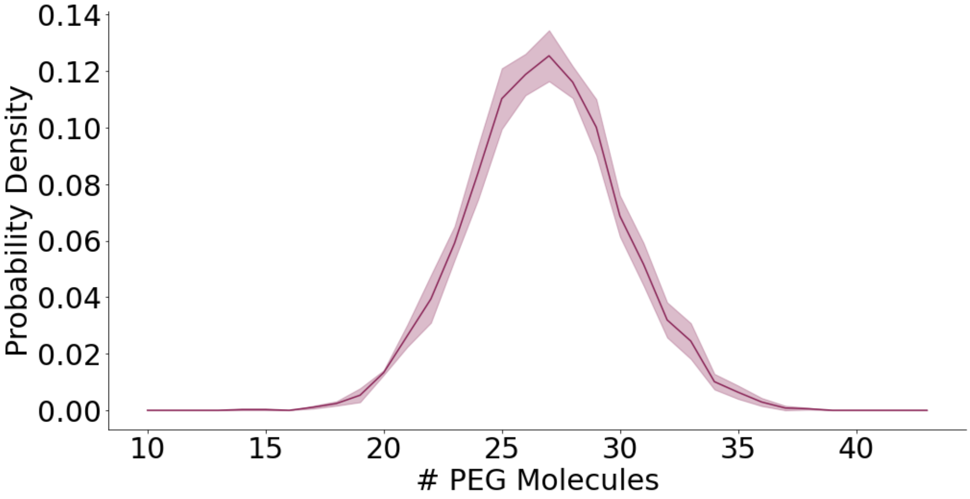


**Figure S13: Number of PEG molecules in contact with mCherry.** The probability distribution of the number of PEG molecules that are in contact with mCherry. Shading indicates standard error of the mean obtained by treating each trajectory as an independent measurement.

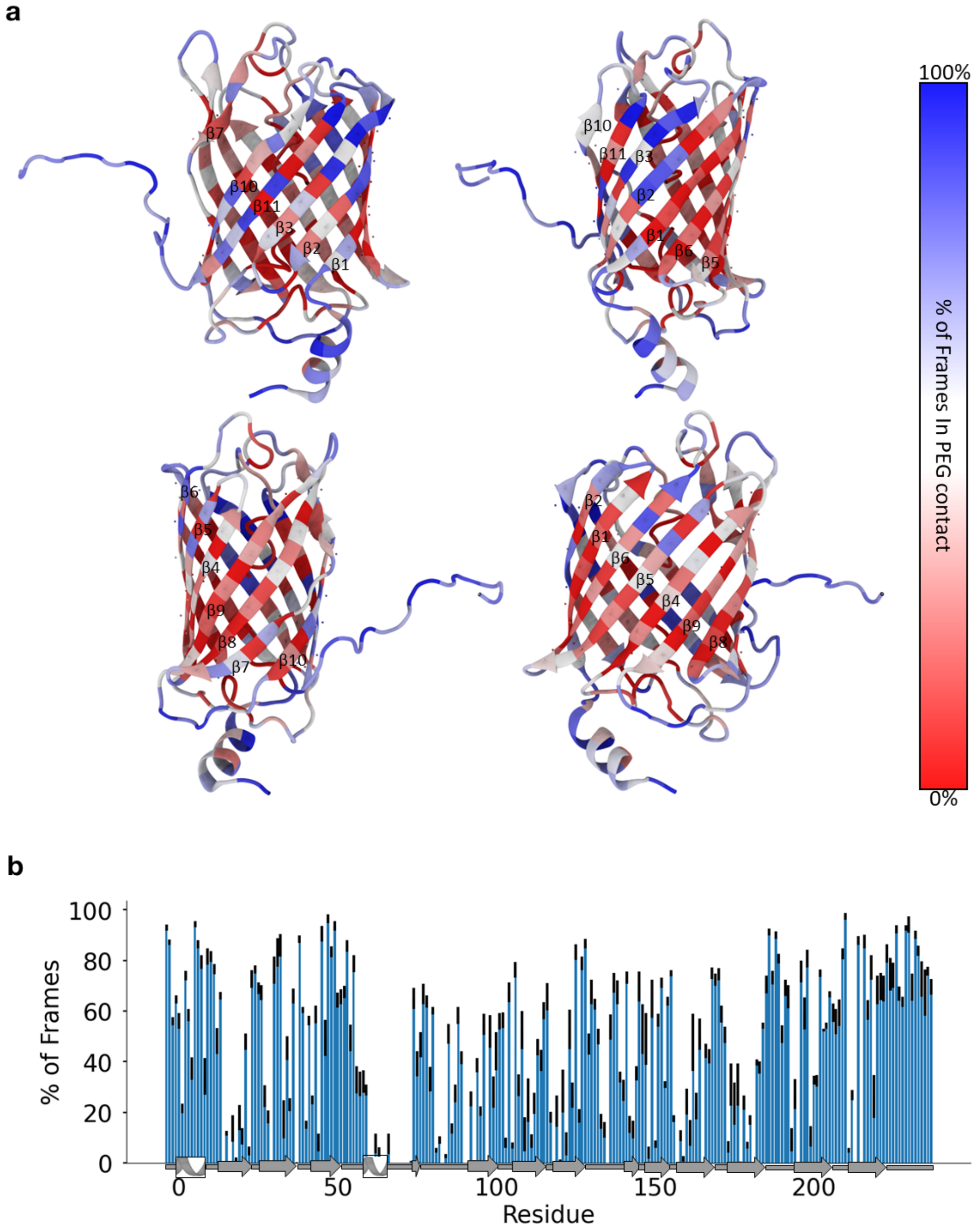


**Figure S14: PEG contact frequency by residue.** (a) mCherry is shown from multiple angles. The protein surface is coloured by the probability of each residue being in contact with PEG. Blue indicates a higher frequency of contacts with PEG. C$\alpha$ atoms are shown as spheres. Side chains that point inwards and outwards from the $\beta$-barrel can be seen from the position of each C$\alpha$ atom. (b) The percentage of frames that each residue is in contact with a PEG molecule. The secondary structure is shown on the x-axis. Error bars indicate standard error of the mean obtained by treating each trajectory as an independent measurement.

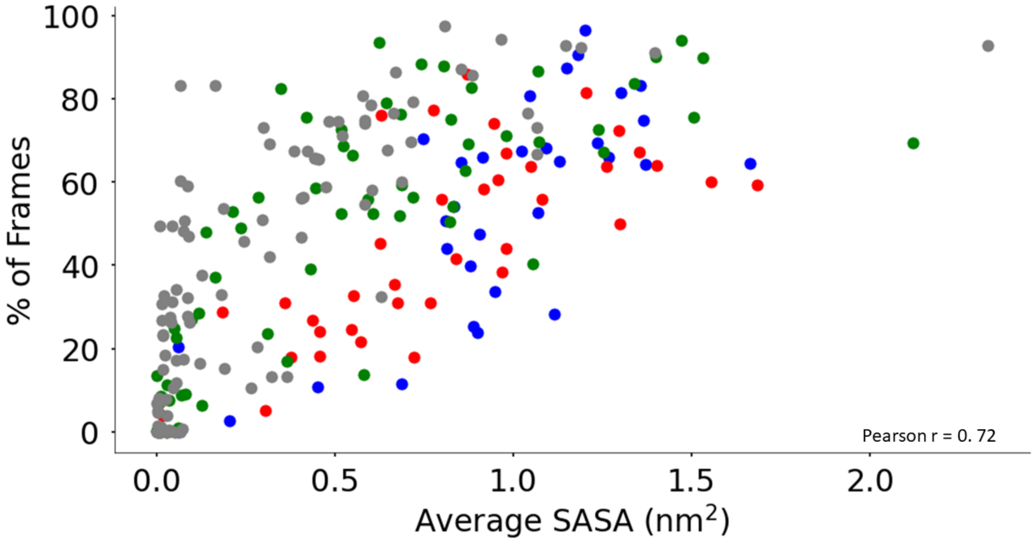


**Figure S15: PEG contact frequency vs. solvent accessibility.** A scatter plot showing the percent of frames that each residue is in contact with PEG vs. the average solvent accessible surface area (SASA). Non-polar residues are shown in grey, negative residues are shown in red, positive residues are shown in blue, and polar residues are shown in green. Pearson r is indicated on the plot.

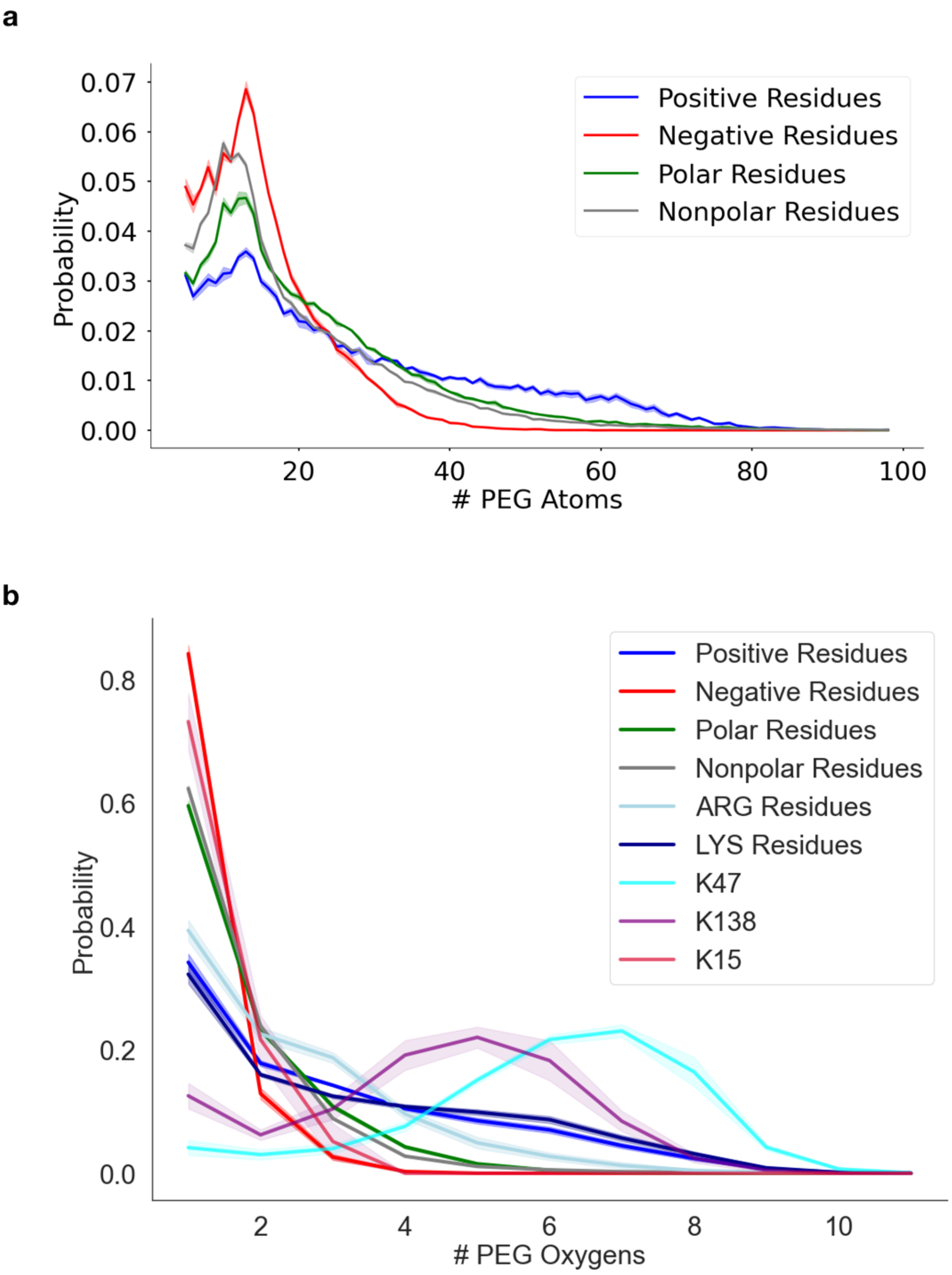


**Figure S16: Contacts with PEG by residue type.** (a) The probability that a residue is in contact with a certain number of PEG atoms, averaged over each residue type. Each PEG molecule contains 199 atoms. Shading indicates the standard error of the mean obtained by treating each trajectory as an independent measurement. (b) The probability that a residue is in contact with a certain number of PEG oxygen atoms, averaged over each residue type, is shown. Shading indicates the standard error of the mean obtained by treating each trajectory as an independent measurement. The distribution for lysine residues K15, K47 and K138 are shown in crimson, cyan and purple, respectively.

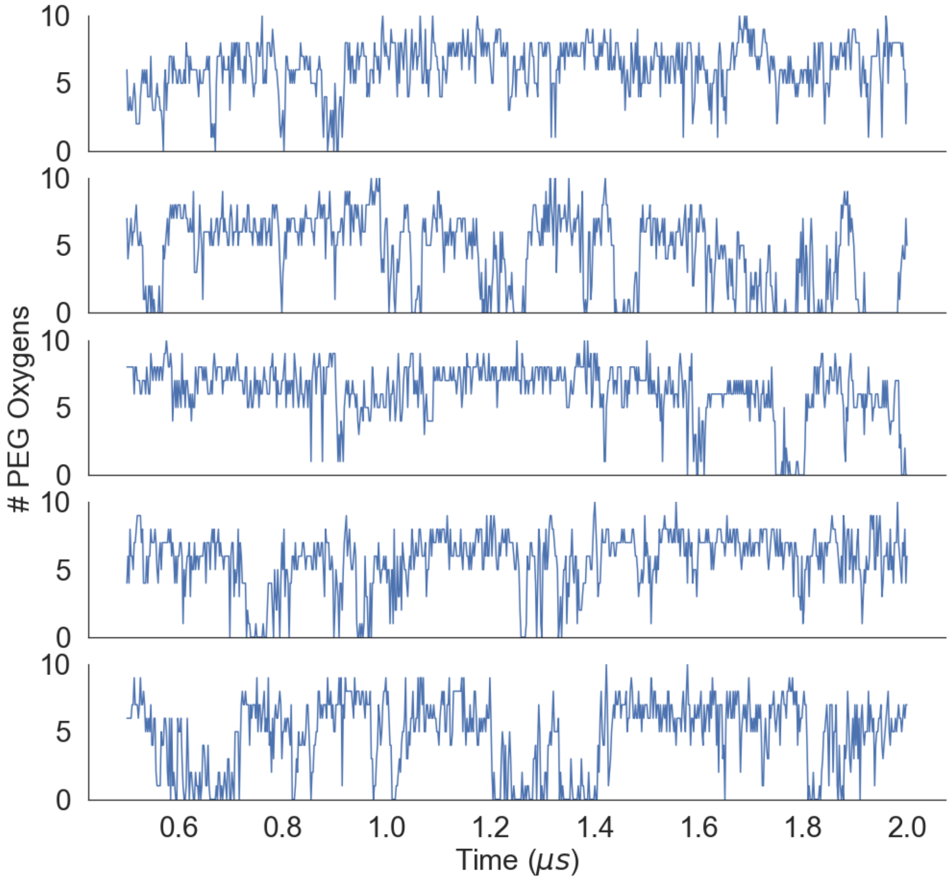


**Figure S17: Time series of K47-PEG interactions.** Time series of the number of PEG oxygen atoms in contact with K47 for each trajectory.

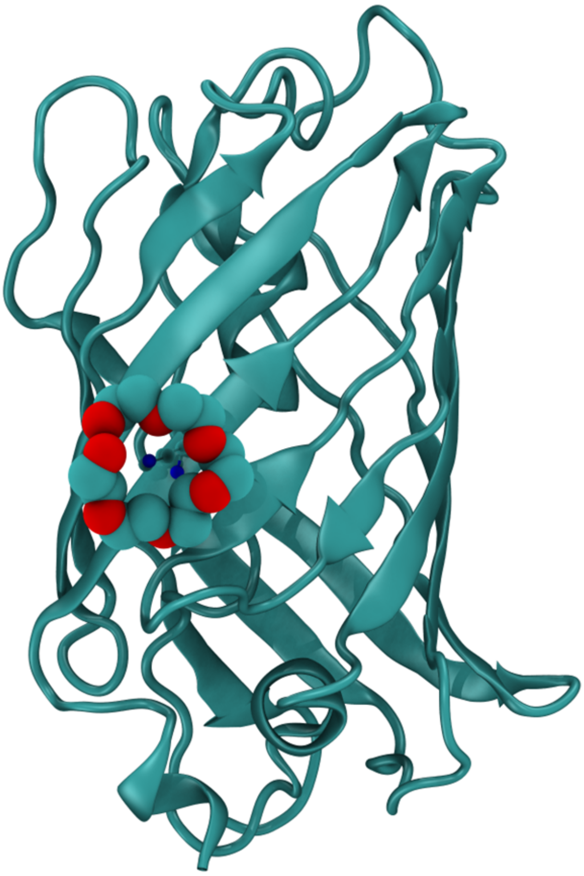

**Figure S18: PEG wrapping observed in mCherry crystal structure** The crystal structure of mCherry from PDB ID: 5FHV[4]. mCherry is shown in cartoon representation, PEG is shown in van der Waals representation, and residue K15 is shown in ball-and-stick representation. While the sidechain of K15 is wrapped by PEG in the crystal structure, this residue is not wrapped significantly in the simulations (Fig. S13b).

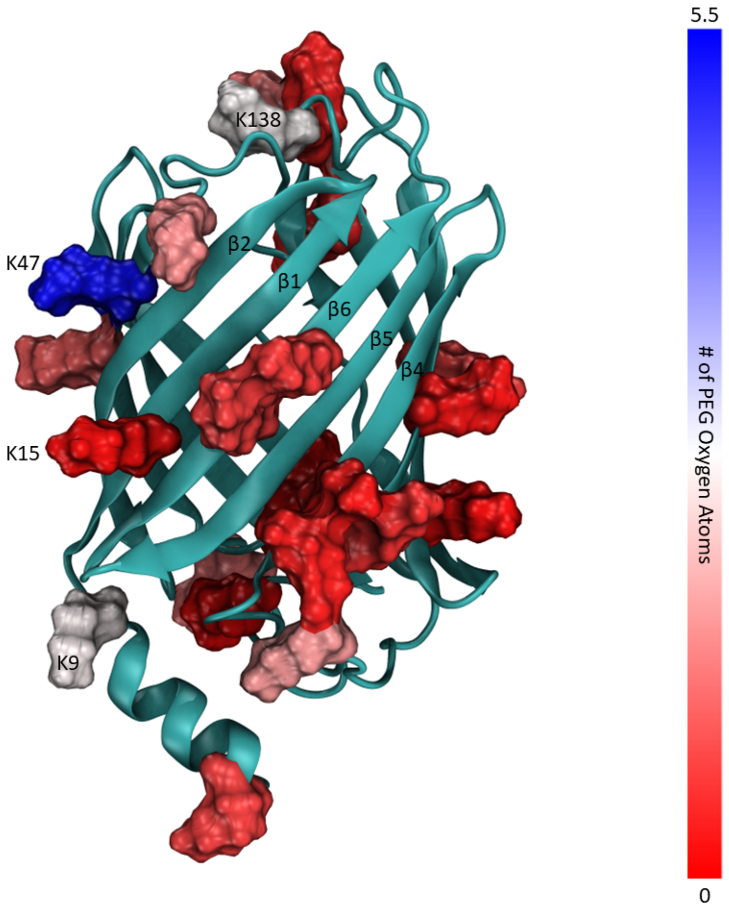


**Figure S19: PEG wrapping varies strongly between lysine residues on the surface of mCherry.** The structure of mCherry is shown in cartoon representation. Lysine residues are shown in surface representation, coloured by the average amount of PEG wrapping observed for each lysine residue in the simulations. The colour scale range is based on the average number of PEG oxygen atoms each residue is in contact with in the simulations. Red indicates little PEG wrapping, white indicates middling PEG wrapping, and blue indicates a high degree of PEG wrapping.

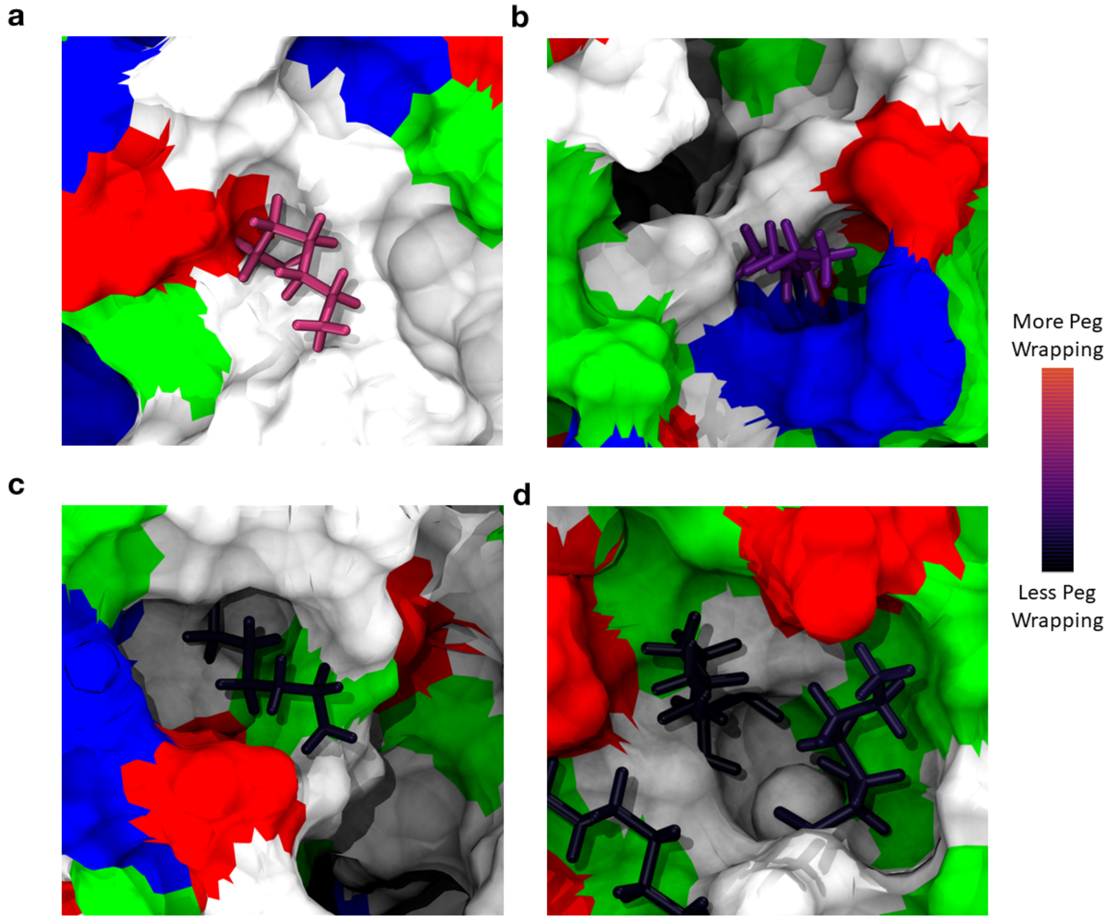


**Figure S20: PEG wrapping in mCherry depends on local environment.** (a-d) Several examples of lysine residues on the surface of mCherry are shown. Lysine residues are shown in licorice representation and coloured by the average number of PEG oxygen atoms with which they form contacts during the simulations. Black indicates fewer PEG oxygen contacts and pink indicates more PEG oxygen contacts. Other residues are shown in surface representation and coloured by residue type. Negatively charged residues are shown in red, positively charged residues are shown in blue, polar residues are shown in green and non-polar residues are shown in white.

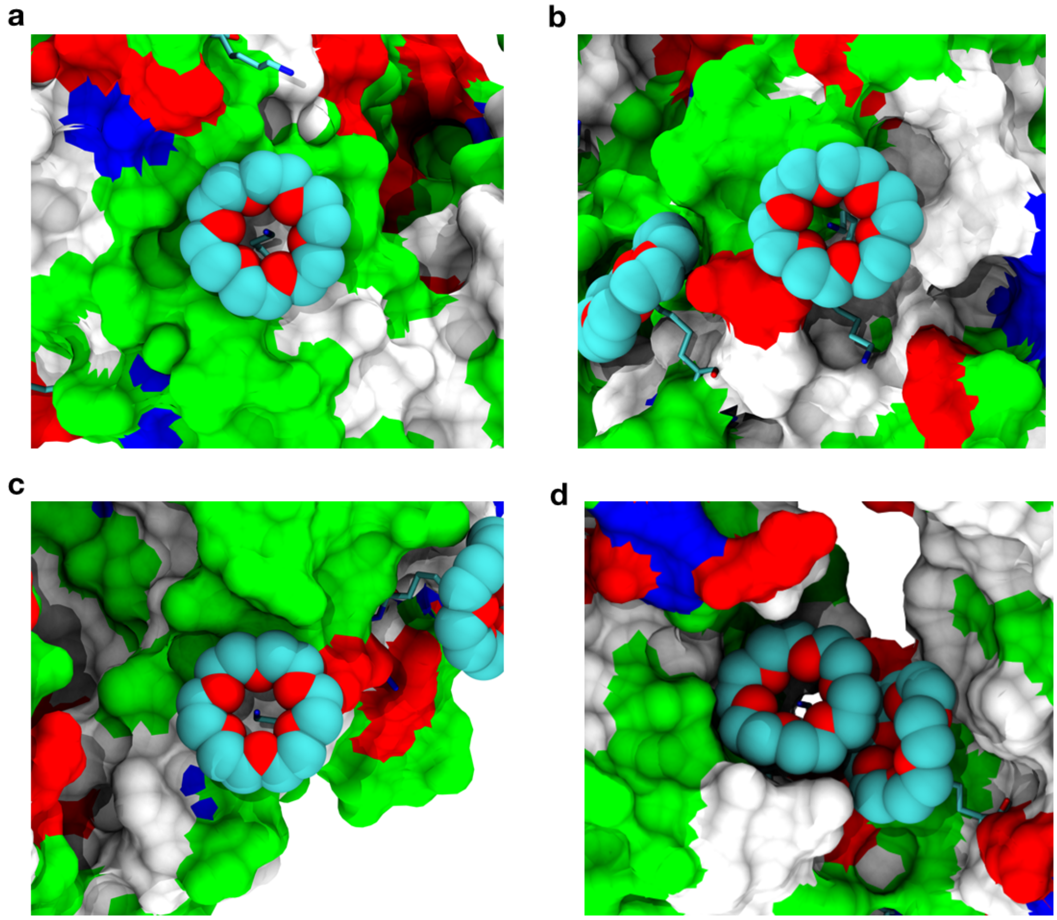


**Figure S21: PEG wrapping in the anti-PEG antibody depends on the local environment.** (a-d) All examples of PEG wrapping from the anti-PEG antibody crystal structure are shown[5] (PDB ID: 6JWC[5]). Lysine residues are shown in licorice representation. PEG molecules are shown in van der Waals representation and coloured by atom type. Other residues are shown in surface representation and coloured by residue type. Negatively charged residues are shown in red, positively charged residues are shown in blue, polar residues are shown in green and non-polar residues are shown in white.

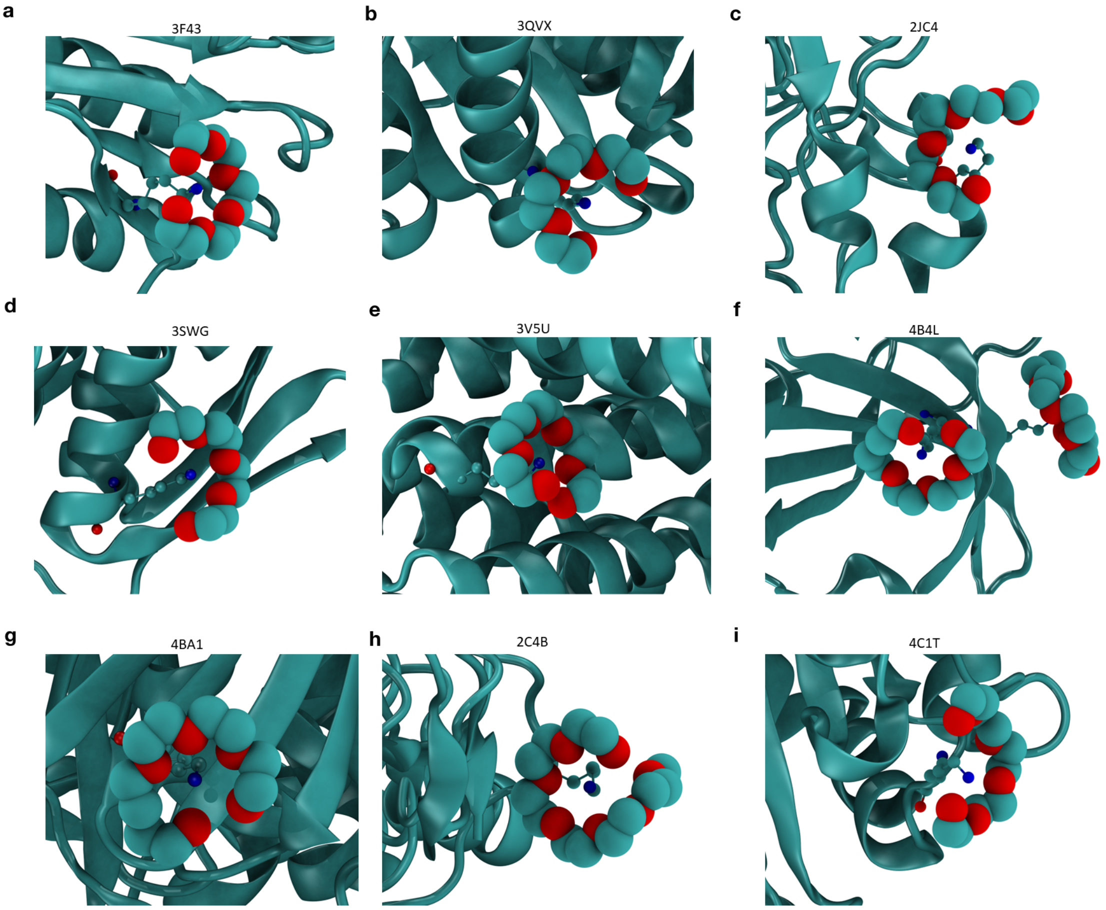


**Figure S22: Examples of PEG wrapping in the PDB.** Several examples of PEG wrapping from structures in the PDB. (a) 3F43[6] (resolution 1.75 Å), putative anti-sigma factor antagonist TM1081 (b) 3QVX[7] (resolution 1.9 Å), Myo-inositol-1-phosphate synthase (Ino1) (c) 2JC4[8] (resolution 1.9 Å), tripeptidase (d) 3SWG[9] (resolution 1.81 Å), UDP-N-acetylglucosamine 1-carboxyvinyltransferase (e) 3V5U[10] (resolution 1.9 Å), Uncharacterized membrane protein MJ0091 (f) 4B4L[11] (resolution 1.75 Å), death associated protein kinase 1 (g) 4BA1[12] (resolution 1.8 Å), probable exosome complex exonuclease 2 (h) 2CB4[13] (resolution 2.5 Å), putative lipoprotein YcdA (i) 4CIT[14] (resolution 1.8 Å), vanadium dependent haloperoxidase. PEG is shown in van der Waals representation. Lysine is shown in ball and stick representation, and the protein is shown in cartoon representation in all panels.

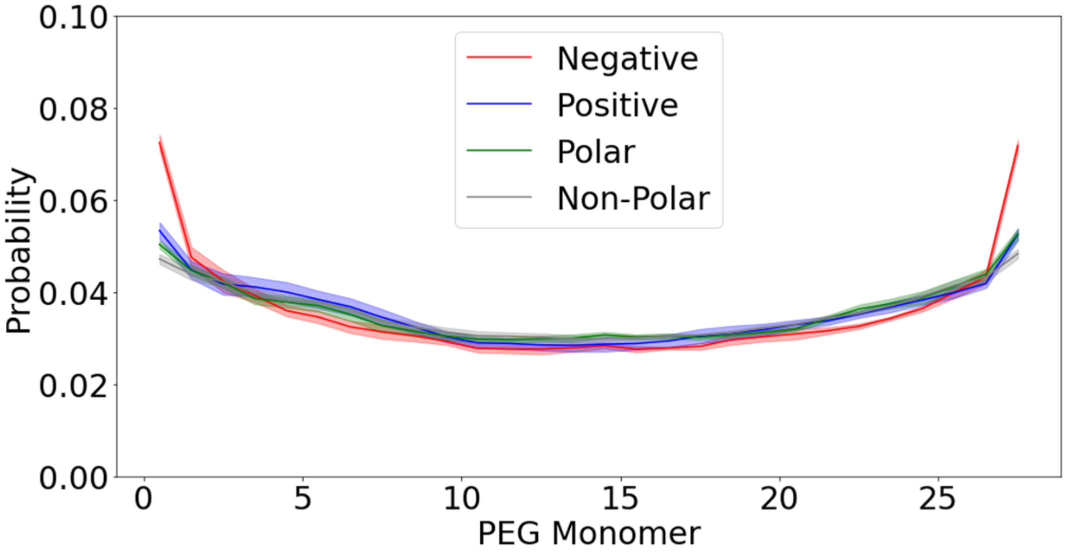


**Figure S23: Contacts between each monomeric unit of PEG and mCherry.** The probability that each of the 28 monomers in the PEG polymer is in contact with protein, separated by residue type. Only PEG molecules that are in contact with protein are counted. The probability is shown for each residue type: positive, negative, polar, and non-polar. Shaded regions indicate the standard error of the mean obtained by treating each trajectory as independent.

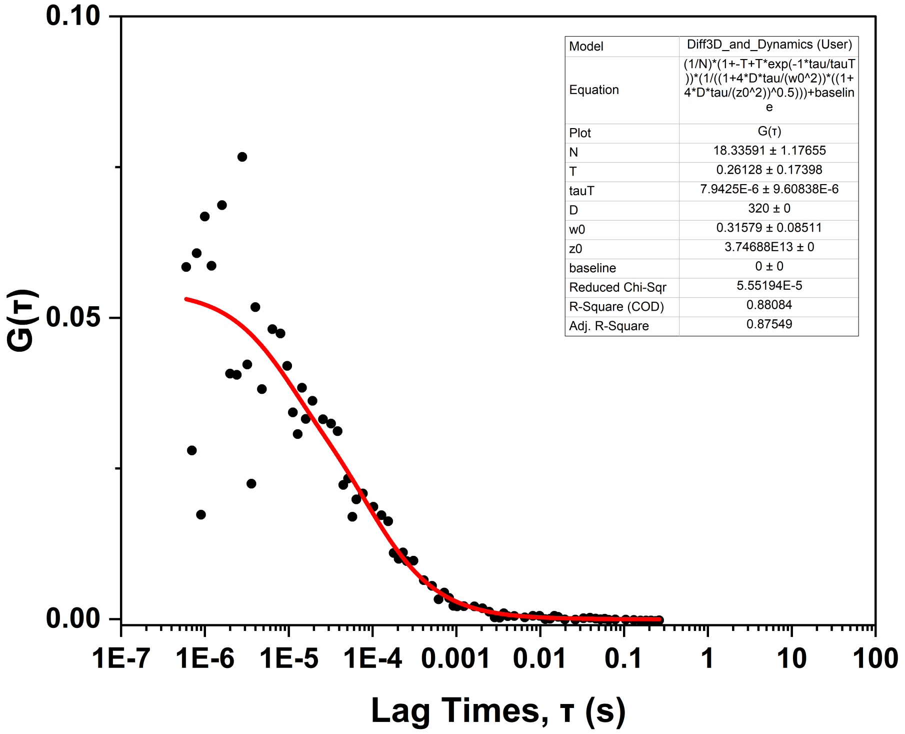


**Figure S24: FCS best fit result for ATTO 532.** The fluorescence autocorrelation curve (black) and the best fit theoretical curve (red) of fluorescence traces of 1 nM ATTO 532 in PBS buffer, using a fixed diffusion coefficient of $\tilde{3}20$ $\mu m^2/s$ [15].

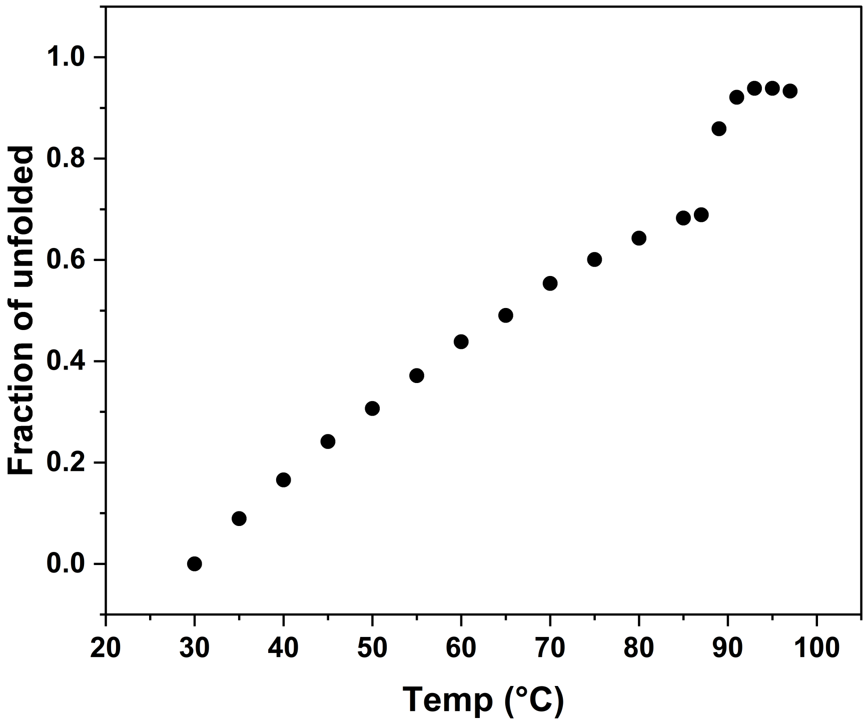


**Figure S25: Thermal stability of mCherry.** Thermal unfolding transition of mCherry recorded by tracking the area below its fluorescence spectrum as a function of temperature using 5°C increments. A gradual decrease in fluorescence and then a global unfolding transition was observed with $T_m$ of $\sim$ 90°C.

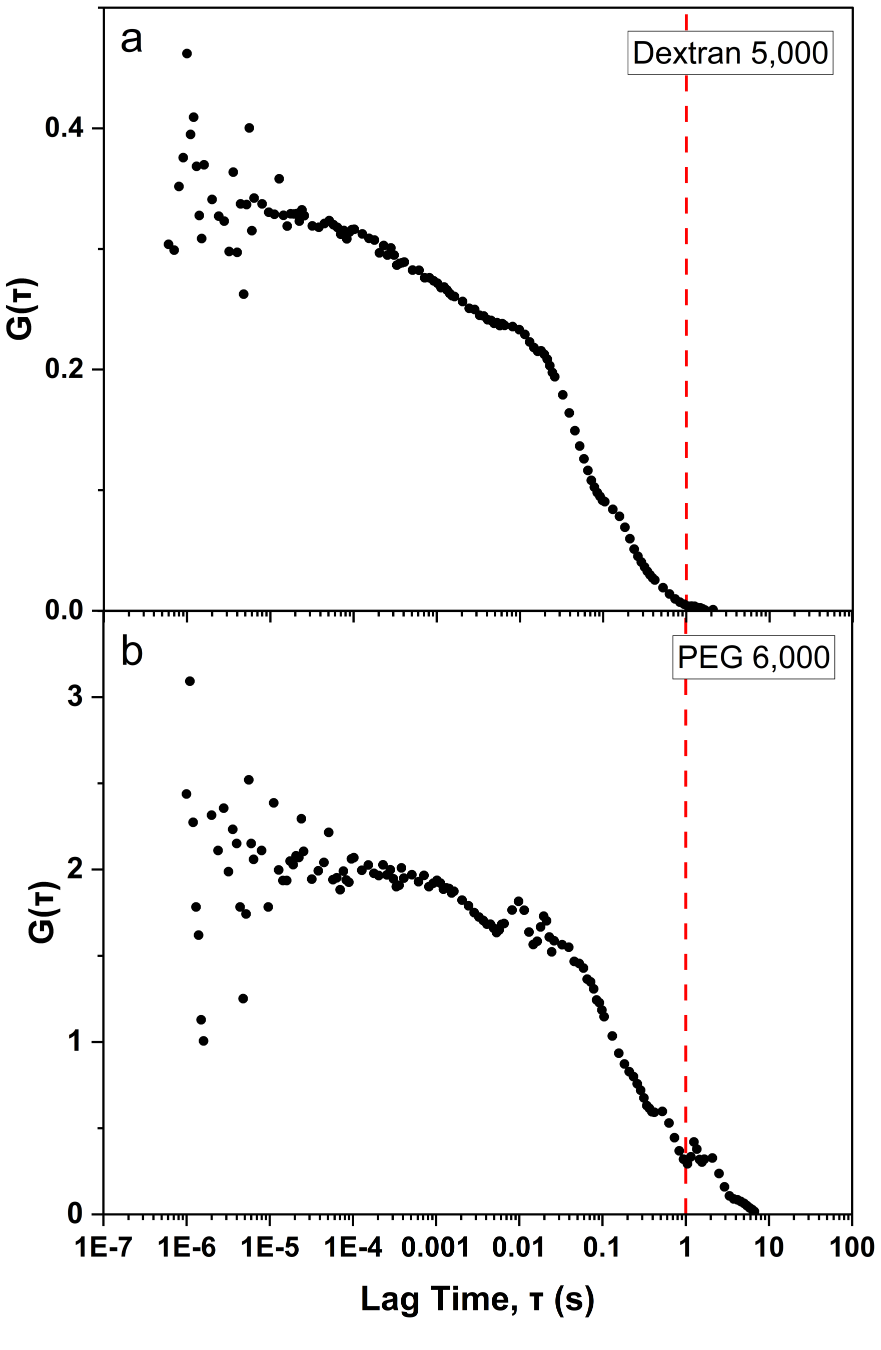


**Figure S26: mCherry aggregates in PEG vs dextran.** The fluorescence autocorrelation curves (a & b, black) of fluorescence traces of 1 nM mCherry in PBS buffer with 40% FVO dextran 5,000 (a) or PEG 6,000 (b). Dashed red line at 1s lag time.

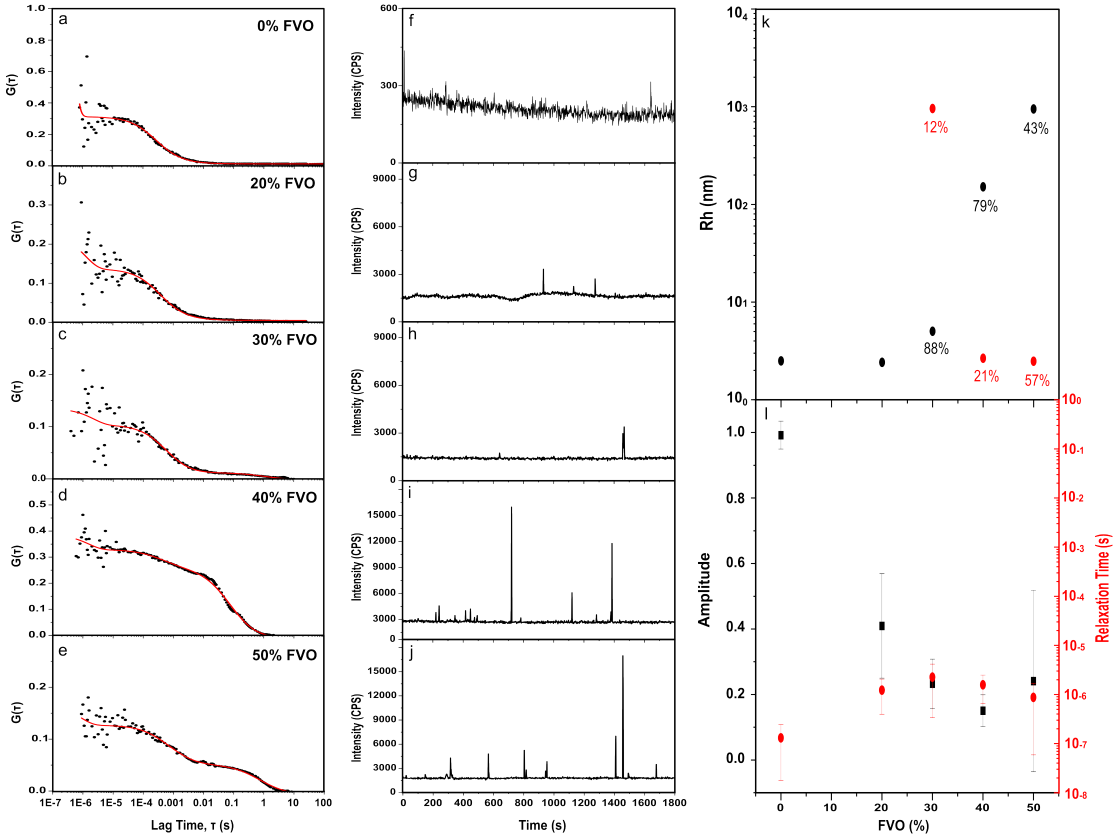


**Figure S27: FCS measurements of mCherry in increasing dextran concentrations.** The fluorescence autocorrelation curves (a-e, black) of fluorescence traces of 1 nM mCherry (f-j, black) in PBS buffer with 0% (a, f), 20% (b, g), 30% (c, h), 40% (d, i) & 50% (e, j) FVO of dextran 5,000, together with the best-fit theoretical curves (a-e, red). Hydrodynamic radius values are calculated from the best-fit diffusion coefficient values (k) using the Stokes-Einstein equation, after factoring out the viscosity of PEG. The hydrodynamic radii are shown per each % FVO dextran for fits to FCS models of single (solely black) or two (black and red) diffusion components. In fits to a model of two diffusion components, the fraction of the diffusion component is provided in the figure panel. The best-fit parameter values are also shown for the rapid fluorescence fluctuation process (l).
